# Using cell yields and qPCR to estimate biotic contribution to 1,1,1-trichloroethane dechlorination at a field site treated with granular zero valent iron and guar gum

**DOI:** 10.1101/583229

**Authors:** M. Ivy Yang, Michael Previdsa, Elizabeth A. Edwards, Brent E. Sleep

**Affiliations:** Department of Civil & Mineral Engineering, University of Toronto, Toronto, M5S 1A4, Canada; Department of Chemical Engineering and Applied Chemistry, University of Toronto, Toronto, M5S 3E5, Canada

**Keywords:** Reductive Dehalogenases, *Dehalobacter*, Groundwater Remediation, Chlorinated Ethanes, Zero Valent Iron, 16S rRNA, Quantitative PCR

## Abstract

Chlorinated ethanes are environmental pollutants found frequently at many contaminated industrial sites. 1,1,1-Trichloroethane (1,1,1-TCA) can be dechlorinated and detoxified via abiotic transformation or biologically by the action of dechlorinating microorganisms such as *Dehalobacter* (*Dhb*). At a field site, it is challenging to distinguish abiotic vs biotic mechanisms as both processes share common transformation products. In this study, we evaluated using the *Dhb* 16S rRNA gene and specific reductive dehalogenase genes as biomarkers for 1,1,1-TCA and 1,1-dichloroethane (1,1-DCA) dechlorination. We analyzed samples from laboratory groundwater microcosms and from an industrial site where a mixture of granular zero valent iron (ZVI) and guar gum was injected for 1,1,1-TCA remediation. Abiotic and biotic transformation products were monitored and the changes in dechlorinating organisms were tracked using quantitative PCR (qPCR) with primers targeting the *Dhb* 16S rRNA gene and two functional genes *cfrA* and *dcrA* encoding enzymes that dechlorinate 1,1,1-TCA to 1,1-DCA and 1,1-DCA to chloroethane (CA), respectively. The abundance of the *cfrA*- and *dcrA*-like genes confirmed that the two dechlorination steps were carried out by two distinct *Dhb* populations at the site. Using cell yields established in laboratory microcosms along with measured abundances of the *Dhb* 16S rRNA gene in site samples, biotic and abiotic transformation of 1,1,1-TCA at the site was estimated. The biomarkers used in this study proved useful for tracking biodechlorination of 1,1,1-TCA and 1,1-DCA where both abiotic (e.g. with ZVI) and biotic processes co-occur.

## 1. Introduction

Chlorinated aliphatic hydrocarbons, such as chlorinated ethanes and ethenes, have been widely used as metal degreasers and cleaning agents, solvents and adhesives, and reagents in various industries (Tobiszewski and Namieśnik, 2012). Of these compounds, 1,1,1-trichloroethane (1,1,1-TCA) was identified as an ozone-depleting compound under the Montreal Protocol of 1987, and its use was phased out starting in 1989 (Gu et al., 2011). Nevertheless, 1,1,1-TCA is still a common organic pollutant in contaminated soil and groundwater at hazardous waste sites, with 1,1,1-TCA found at more than 393 of the 1315 active sites on the National Priorities List (US EPA database as of 2012). Exposure to 1,1,1-TCA may lead to damage to the central nervous system, heart and liver (ATSDR, 2006). The EPA has set the maximum contaminant level of 1,1,1-TCA in drinking water at 200 µg/L (EPA, 1995).

In groundwater, 1,1,1-TCA is prone to abiotic transformation to 1,1-dichloroethene (1,1-DCE) via elimination, or to 1,1-dichloroethane (1,1-DCA) via reduction catalyzed either by naturally occurring metals or metals added in engineered systems (Scheutz et al., 2011; Palau et al., 2016). In particular, the addition of zero valent iron (ZVI) has been tested at lab and pilot scales and implemented at various sites to reduce chlorinated compounds including 1,1,1-TCA. At these sites 1,1-DCA was often found as the major daughter product of 1,1,1-TCA and ethane as a minor by-product (Fennell and Roberts, 1998; Lookman et al., 2004; Velimirovic et al., 2014; Wu et al., 2014).

Biotransformation of 1,1,1-TCA occurs in anaerobic environments via reductive dechlorination (Fam et al., 2012). *Dehalobacter* (*Dhb*) sp. TCA1 was the first pure culture of an organohalide respiring bacterium that coupled growth to the sequential dechlorination of 1,1,1-TCA to 1,1-DCA and ultimately to chloroethane (CA) (Sun et al., 2002). A few other strains, namely *Dhb* sp. strain CF and strain DCA (Tang and Edwards, 2013B), *Dhb* sp. strain UNSWDHB (Wong et al., 2016), *Dhb* sp. strain THM1 (Zhao et al., 2017), and *Desulfitobacterium* strain PR (Ding et al., 2014), have since been described or isolated with the ability to dechlorinate 1,1,1-TCA and/or 1,1-DCA. All of these strains use either hydrogen or formate as electron donor. Five reductive dehalogenases (RDases) catalyzing these specific dechlorination reactions in these bacteria have also been identified (i.e., CtrA, TmrA, DcrA, CfrA and ThmA). These proteins form a clade of highly similar sequences belonging to Ortholog Group 46 as defined by Hug et al.(2013) and updated by Molenda et al. (2020). Interestingly, while the proteins share greater than 95% amino acid identity, they do have distinct substrate preferences. For example, DcrA catalyzes dechlorination of 1,1-DCA to CA, but is not active on 1,1,1-TCA or chloroform, while CfrA and ThmA are active on chloroform and 1,1,1-TCA but not on 1,1-DCA (Tang and Edwards, 2013; Tang and Edwards, 2013B; Zhao et al., 2017). CtrA from *Desulfitobacterium* strain PR and TmrA from *Dhb* UNSWDHB are reported to be active on all three substrates to varying degrees (Ding et al., 2014; Wong et al., 2016).

Chlorinated ethane concentrations are frequently monitored to evaluate 1,1,1-TCA treatment efficiency at remediation sites, but concentration data are easily affected by site operations, making interpretation of data difficult, particularly in trying to distinguish between biotransformation and abiotic removal. Gene-based biomarkers applied to field samples have the potential for monitoring biological 1,1,1-TCA removal (Scheutz et al., 2011). However, only a limited number of field studies reported using quantitative PCR (qPCR) with primers to measure the 16S rRNA gene of *Dehalobacter* dechlorinating 1,1,1-TCA at remediation sites (Postiglione et al., 2006; Duchesneau et al, 2007; Damgaard et al., 2013). Moreover, to date, primers targeting functional genes from organisms dechlorinating 1,1,1-TCA have only been applied to microcosm and culture samples, but not to field samples in published work. Without specific biomarkers, estimating the fraction of 1,1,1-TCA removed by biodegradation remains challenging.

In this study, we hypothesized that the change in chlorinated ethane concentration caused by biodegradation would be closely reflected in the abundance of substrate-specific dehalogenase genes, and that the fraction of 1,1,1-TCA removed by reductive dehalogenation at a contaminated site could be estimated based on cells yields and the abundance of these biomarker genes. To test this hypothesis, we worked in collaboration with consultants from Pinchin Ltd. on a site contaminated with 1,1,1-TCA. The *in situ* remedy selected for the site was injection of granular ZVI mixed with guar gum. The objectives of this research were to evaluate the use of the *Dhb* 16S rRNA gene and the *cfrA*- and *dcrA*-like dehalogenase genes as biomarkers for 1,1,1-TCA and 1,1-DCA biodechlorination, and ultimately to estimate the fraction of 1,1,1-TCA biotransformed at the site. Additionally, the impact of guar gum and ZVI addition on the biodegradation of 1,1,1-TCA was assessed. This project consisted of the analysis of site samples as well as a laboratory groundwater microcosm study. Site samples were collected to measure chlorinated ethane concentrations and gene abundances using qPCR during the on-going remediation. Microcosms prepared with site groundwater were used track both abiotic and biotic processes under more controlled conditions, and to evaluate the specificity of the substrate-specific primer sets, and to estimate approximate cell yields applicable to the corresponding site samples.

## 2. Materials and Methods

### 2.1 Site Description

The Site (Supplementary Fig. S1) was a former adhesive manufacturing plant located in Southern Ontario and operated between 1950 and 1991. A spill of approximately five hundred litres of 1,1,1-TCA in the late 1980s was the only recorded release at the site. Historical contaminant concentrations are presented in Supplementary Table S1. Prior to the guar gum-ZVI injection, 1,1,1-TCA was the dominant contaminant at the site, with concentrations approximately ten times greater than other compounds. The stratigraphy of the site is provided in Supplementary Text S1.

### 2.2 Granular ZVI Particles, Guar Gum, and On-Site Guar Gum-ZVI Injection

In late September 2015, the mixture of guar gum and ZVI was injected mainly along the property boundary and in the plume area (Supplementary Fig. S1). Guar gum is a polysaccharide mixture extracted from guar beans and is predominantly composed of galactose and mannose. One week after guar gum-ZVI injection, portions of the site were excavated and back-filled with ZVI alone. The details of ZVI, guar gum matrix, fracturing pressure, distance and intervals of injections, and the amounts of guar gum and ZVI injected and soil excavation are explained in Supplementary Table S2.

### 2.3 Groundwater Monitoring and Sample Collection

Between September 2015 and June 2016, groundwater samples were collected from three wells in the plume area, SW5, SW6 and MW202 (Supplementary Fig. S1). In the field study presented in this paper, we defined Day 0 as the time when the first groundwater sample was collected prior to ZVI injection. All wells were developed and purged prior to sampling. Upon groundwater recovery, samples were collected using dedicated inertial pumps. Groundwater samples for chlorinated organic compound analysis were stored in 44 mL glass vials with screw caps with Teflon liners and zero headspace, and preserved with 0.2 g sodium bisulphate tablets. Samples for microbial analysis were collected in 250 mL sterilized plastic bottles containing sodium thiosulphate with zero headspace. All samples were placed on ice immediately after sampling and arrived at our lab at the University of Toronto on the same day.

### 2.4 Groundwater Microcosm Setup and Sampling

To evaluate *in situ* microbial activity with and without ZVI added, groundwater collected on Day 202 from SW6 was used to set up microcosms. The groundwater (2 × 2 L in medium bottles) was purged with 80%/20% N_2_-CO_2_ gas mix for 1 hour to completely remove oxygen and residual volatile compounds. In a disposable glovebag (Atmosbag, Sigma-Aldrich, Missouri, US) filled with 80%/20% N_2_-CO_2_ gas mix, 150 mL of purged groundwater was dispensed into each sterile 250 mL glass bottle, and one gram ZVI (same type used at the site) was added to the bottles designated with ZVI addition. Mininert septa screw caps were sterilized by wiping with ethanol. All bottles were then capped with the sterile Mininert caps before leaving the glovebag. No other inoculum, electron donor or carbon source was added, unless specified.

As detailed in Supplementary Table S3, six treatments were established, including: Sterile (i.e., negative control by autoclaving); Sterile + ZVI; Active without ZVI; Active + ZVI; Active without TCA and ZVI; and bioaugmented with ACT3 (ACT3 + ZVI) where ACT3 refers to a previously described enrichment culture that dechlorinated 1,1,1-TCA via 1,1-DCA to CA (Tang and Edwards, 2013B). Neat 1,1,1-TCA (∼4.25 μL per bottle) was added for an initial liquid concentration of approximately ∼0.2 mmol/L (0.036 mmol per bottle). In the groundwater microcosm experiment, Day 0 refers to the day when the batch assays were set up and 1,1,1-TCA was added to the bottles. ACT3 culture (0.1 mL) was anaerobically added to the ACT3 + ZVI set on Day 2 inside a lab glovebox, yielding an additional ∼1×10^9^ copies of the *Dhb* 16S rRNA gene /L in these ACT3 + ZVI bottles. In most of the microcosm bottles, we did not add any additional electron donor, as the purged groundwater contained approximately 800 mg/L COD from residual guar gum and its degradation products. The only exceptions were bottle #8 in the Active set without ZVI and bottle #21 in the Active + ZVI set. Ethanol and lactate were added to these two bottles on Day 149 at five times electron equivalent excess to the remaining 1,1-DCA (Supplementary Table S3). An extra bottle containing the same amount of groundwater without any 1,1,1-TCA or ZVI was also included in the assay to evaluate methane production from the groundwater COD. All bottles were incubated statically, upside-down and covered with black cloth in an anaerobic glovebox (Coy Laboratory Products Inc, Michigan, US) filled with 5%-95% H_2_-N_2_ gas mix at room temperature.

### 2.5 DNA Extraction and Microbial Analysis

Groundwater samples collected from the site for microbial analysis were packed with ice and shipped to our lab at the University of Toronto, and filtered within 24 hours after sampling through sterile 0.22 μm Sterivex filters using a vacuum pump (Ritalahti et al., 2009). The filters were frozen at −80 °C immediately after filtration. Liquid samples from lab microcosms (1.6 mL) were collected using sterile plastic syringes and needles, and transferred to 2 mL anaerobic screw-cap vials inside the glovebox. Samples were centrifuged at 13,000×g at 4 °C for 15 minutes. The supernatant was removed and the vials with pellets were stored at −80 °C until DNA extraction.

DNA extraction was conducted using the Mobio® PowerSoil DNA isolation kit (MoBio Laboratories Inc., Carlsbad, CA). For groundwater samples, filter membranes were removed from the plastic casings, and added to the bead beating tubes. For each microcosm sample, the beads and the solution in one bead beating tube were carefully transferred to the screw-cap vial containing the thawed cell pellet, and vortexed until the cell pellet was completely re-suspended. Subsequent DNA extraction steps were performed following manufacturer instruction. A NanoDrop ND-1000 spectrophotometer was used to measure DNA concentration.

Quantitative PCR (qPCR) was used to enumerate specific phylogenetic and functional genes in the DNA samples. Primers *Dhb*477f ((5’-GATTGACGGTACCTAACGAGG-3’) /*Dhb*647r (5’-TACAGTTTCCAATGCTTTACGG-3’) were used to quantify the abundance of the *Dhb* 16S rRNA gene copies (Grostern and Edwards, 2006). Primers *cfrA*413f (5’-CCCGAACCTCTAGCACTTGTAG -3’) /*cfrA*531r (5’-ACGGCAAAGCTTGCACGA -3’) and *dcrA*424f (5’-AGCACTCAGAGAGCGTTTTGC -3’) /*dcrA*533r (5’-CAACGGCCCAGCTTGCAT -3’) were adapted to track two reductive dehalogenase genes *cfrA* and *dcrA* encoding the proteins catalyzing dechlorination of 1,1,1-TCA and 1,1-DCA respectively (Tang and Edwards, 2013B). The annealing temperature was set at 62.5 °C for the *Dhb* 16S primer or at 60 °C for the *cfrA* and *dcrA* primer sets. Detailed recipe and information of the qPCR reactions and detection are presented in Supplementary Text S2 and Fig. S2.

The DNA extracted from site and microcosm samples was also submitted to McGill University and Genome Quebec Innovation Center for amplicon sequencing with the universal 16S rRNA gene primers 926f (5’-AAACTYAAAKGAATWGRCGG-3’) and 1392r (5’-ACGGGCGGTGWGTRC-3’) (Qiao et al., 2020). The amplicon sequencing data were analyzed using the QIIME2 pipeline with the Dada2 package (Bolyen et al., 2019, details in Supplementary Text S3).

To obtain nearly complete sequences of the reductive dehalogenase genes from our samples, the DNA extracted from the bottle with the fastest degradation of 1,1,1-TCA (bottle #9 on Day 49) was amplified with primers designed by Tang and Edwards (2013B) termed *rdhA*23f (5’-AAGAGATTGTAhGAAGCAGCGG-3’) and *rdhA*1383r (5’-CTTAGTAAATGGGCAAGCAGC-3’) targeting the RDase genes (the original numbering is a little mis-leading as the forward primer is actually a little outside the coding region, as shown in Supplementary Text S4). The PCR program, the primer sequences, the cloning procedures in *E. coli* and the procedures to extract plasmid DNA are described in Tang and Edwards (2013B). Clones were submitted to The Centre for Applied Genomics (Toronto), and each was sequenced four times in both the forward and reverse directions with the M13f and M13r primers. The forward and reverse reads were assembled using Geneious and resulting contigs aligned with the five known gene sequences encoding 1,1,1-TCA/1,1-DCA dehalogenases: *dcrA* (JX282330.1), *cfrA* (JX282329.1), *ctrA* (KF155162.1), *tmr*A (NZ_AUUR01000010 REGION: 19936..21303) and *thmA* (KX344907.1).

### 2.6 Chemical Analysis of Groundwater and Microcosm Samples

Chlorinated ethane concentrations in field samples were stored at 4 °C and were measured within 48 hours after the samples arrived at the University. Microcosm were first shaken and then sampled using gas-tight glass syringes and sterile needles inside the glovebox, and analyzed immediately. Aliquots (0.5 – 1 mL) from groundwater or microcosm samples were added to acidified MilliQ water to a total liquid volume of 6 mL in 11 mL crimped-cap headspace autosampler vials and placed in the Agilent G1888 Headspace Sampler. Headspace was analyzed using the Agilent 7890 gas chromatograph (GC) equipped with the GS-Q column and detected using a flame ionization detector (FID). External standards, purchased from Sigma, were first diluted with methanol, and then in acidified MilliQ water for calibration. A modified EPA 8021 method was used as per Kocur et al. (2015). Several groundwater samples were also analyzed for total COD according to the EPA Method 410.4 (USEPA, 1993).

### 2.7 Accession Numbers

Raw amplicon sequences were submitted to the NCBI Sequence Read Archive (SRA, ID PRJNA634323), with sample accession numbers from SAMN14988330 to SAMN14988369 (details in Supplementary Text S3). The cloned sequences of the nearly complete RDase genes were deposited to GenBank with accession numbers MT536772 for Clone 1, MT536773 for Clone 2 and MT536774 for Clone 3 (sequences also provided in Supplementary Text S4).

## 3. Results and Discussion

We analyzed chlorinated solvent concentrations and gene abundances in both microcosm and groundwater samples to investigate transformation of 1,1,1-TCA at a field site undergoing treatment with guar gum stabilized ZVI. The microcosms provided a controlled experiment to distinguish biodegradation from abiotic transformation, to obtain a correlation between depletion of chlorinated ethanes and changes in biomarker abundance, and to aid in interpreting biodegradation at the site. Microcosm results are presented first, followed by field data, and ending with an analysis of microbial communities in both lab and field samples. The contributions of abiotic and biotic processes are also discussed.

### 3.1 Dechlorination of 1,1,1-TCA in Groundwater Microcosms

Six treatments were evaluated in microcosms prepared with site groundwater only (Supplementary Table S3). To evaluate the accuracy of our measured concentration data, the sum of the moles of all two-carbon compounds (C2-compounds) was tracked over time considering all possible products of 1,1,1-TCA dechlorination. These included 1,1-DCA, 1,1-DCE, cis-DCE, trans-DCE, 1,2-DCA, CA, ethane and ethene. The sum of moles of all C2-compounds was conserved in the Sterile Control microcosms without ZVI, in the Active set without ZVI, and in the set with ACT3 culture plus ZVI (ACT3 + ZVI), confirming that all major transformation products were being monitored (Supplementary Fig. S3) in this cases. In the Active + ZVI and Sterile + ZVI sets, measured degradation products accounted for only 70-80% of substrate (1,1,1-TCA) loss, indicating that some products were not being measured in these bottes (discussed further below). Methane production was also monitored and compared to the original COD concentration measured at Day 0 in microcosms (Supplementary Table S4A). In the absence of 1,1,1-TCA and ZVI, methane production accounted for approximately 90% of the original COD present in the bottle, indicating that although no extra electron donor was added to the microcosm bottles, residual guar-gum and transformation products in the groundwater were sufficient to drive methanogenesis (and dechlorination in the bottles fed with 1,1,1-TCA). Less methane was produced in bottles fed with 1,1,1-TCA reflecting the well-documented inhibition of methanogenesis (Grostern and Edwards, 2006B). Mole balances were closed in most cases establishing the accuracy of our analytical data.

Dechlorination of 1,1,1-TCA to 1,1-DCA occurred in all cases except for the Sterile Control bottles without ZVI (Supplementary Fig. S4A). However, the extent of dechlorination, the lag time, and the end-products varied considerably, even within replicates for a given treatment (Fig. 1 & Fig. S6). For example, dechlorination to CA was observed in bottles #7 and #9 from the Active without ZVI treatment, but not in bottle #8 from the same set until it was amended with electron donor after Day 150 (Fig. 1). Dechlorination of 1,1-DCA to CA was also seen in bottle #18 of the Active + ZVI treatment, but not in bottle #21 until it was amended with electron donor after Day 150 (Fig. S6). Dechlorination to CA was also observed in the ACT3 + ZVI treatment (Fig. S5). In contrast, in the Sterile + ZVI bottles (Supplementary Fig. S4B), dechlorination of 1,1,1-TCA stopped at 1,1-DCA, consistent with previous studies reporting that 1,1-DCA is not reduced abiotically by ZVI (Cwiertny et al., 2006; Wu et al., 2014). In all bottles containing ZVI, whether (biologically) active or not, dechlorination of 1,1,1-TCA to 1,1-DCA was relatively rapid (i.e., completed in less than 50 days). In bottles without ZVI, lag times ranged from 7 to 21 days before initiation of dechlorination, although once initiated, dechlorination rates were similar in all these bottles. Ethane and ethene were only detected in the ZVI-containing bottles (Table S4B), and these minor products appeared as early as Day 1, possibly via another abiotic dechlorination pathway bypassing CA, as no CA was detected in the Sterile + ZVI set. Moreover, in the bottles containing ZVI (sterile or not), we observed a 20-30% mole loss over the first two weeks (Supplementary Fig. S3), presumably due to the formation of unidentified products such as 2-butyne or cis-2-butene, as previously reported (Fennelly and Roberts, 1998). In summary, 1,1-DCA was the major final dechlorination product from 1,1,1-TCA with ZVI alone. In the presence of active microorganisms, dechlorination proceeded further to CA. The addition of ZVI reduced the lag time for 1,1,1-TCA dechlorination to 1,1-DCA and generated other by-products that were not identified.

**Fig. 1.**
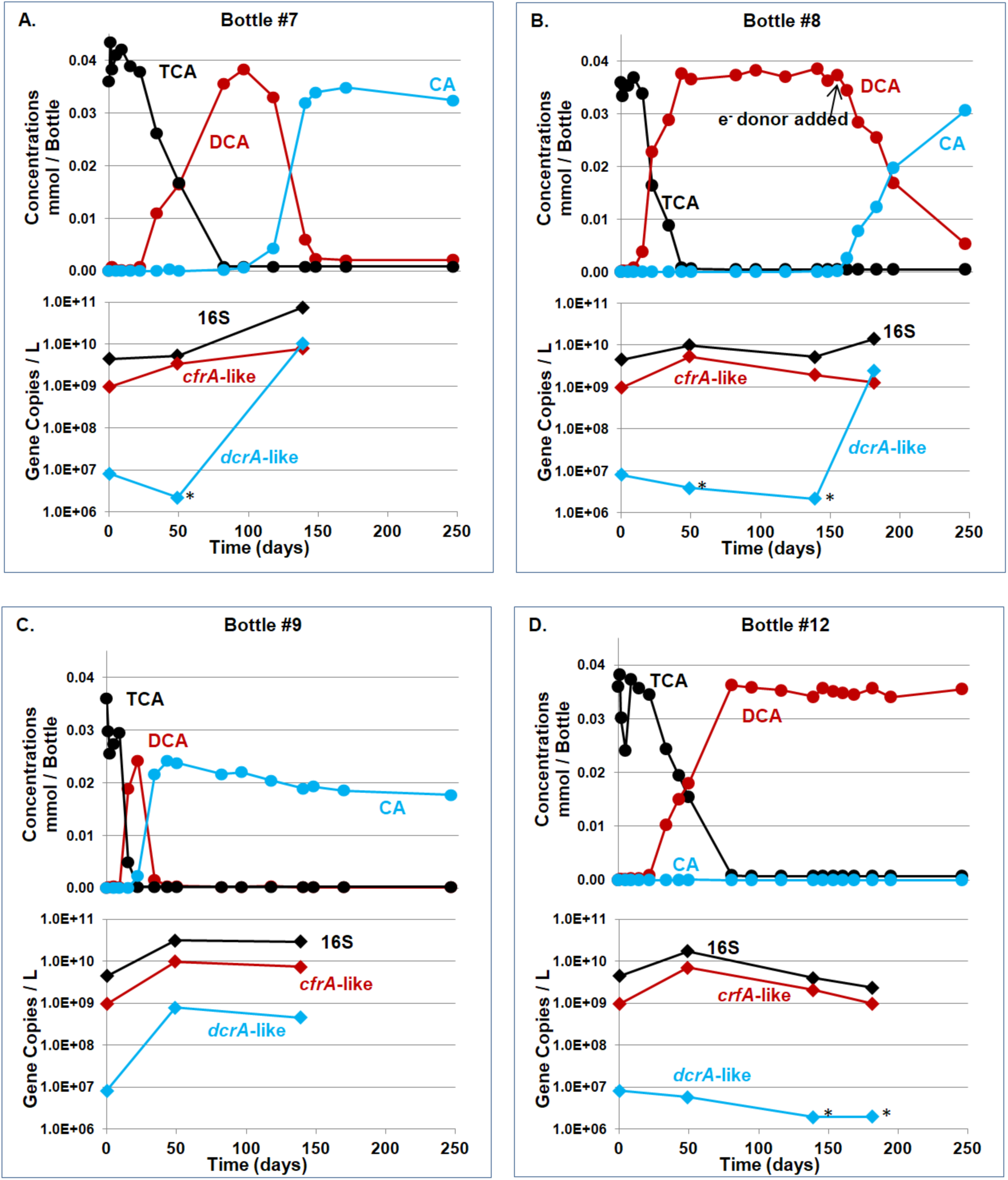
Chlorinated ethane concentrations and gene copy numbers in groundwater microcosms without ZVI. Data from bottles #7, #8, #9 and #12 are presented in Panels A, B, C and D, respectively. In each panel, the top figure shows chlorinated ethane concentrations (in circles, black for 1,1,1-TCA, red for 1,1-DCA, and blue for CA) and the bottom figure displays the qPCR results in copies per L (in diamonds, black for the *Dhb* 16S rRNA gene, red for the *cfrA*-like gene, and blue for the *dcrA*-like gene). Gene copies are the average of three technical replicates, and qPCR data points marked with * are below the lowest detection limit or showed nonspecific amplification.

### 3.2 Evaluation of the Specificity of the Primer Sets Targeting the *cfrA*-like and *dcrA*-like RDase Functional Genes

Certain *Dehalobacter restrictus* (*Dhb*) populations are well-known for catalyzing 1,1,1-TCA and 1,1-DCA dechlorination, therefore we first established that this genus was present in the groundwater, and then focussed on enumerating *Dhb* in the microcosms over time. We also tracked the abundance of the corresponding functional genes *cfrA* and *dcrA* (Tang and Edwards, 2013B). We amplified and cloned the DNA extracted from microcosm bottle #9 on Day 49 to retrieve nearly complete (1231bp) RDase genes. Three clones were successfully sequenced, and their sequences were compared to the five genes encoding RDases known to dechlorinate 1,1,1-TCA and/or 1,1-DCA, namely CtrA, TmrA, ThmA, CfrA and DcrA (Tang and Edwards, 2013B; Ding et al., 2014; Wong et al., 2016; Zhao et al., 2017). Based on sequence alignment and clustering (Fig. S7A and Table S5), Clone 3 was found to be most similar (>99.5%) to the *dcrA* gene (encoding a 1,1-DCA dehalogenase), and Clones 1 and 2 were more similar to the *thmA* gene which encodes an RDase that is active on 1,1,1-TCA but not 1,1-DCA, similar to CfrA (Tang and Edwards, 2013B; Zhao et al., 2017). Next we aligned the cloned sequences to the *cfrA* and *dcrA* primer sequences. As shown in Table 1 (detailed alignment in Fig. S7B). the alignment revealed two clades. The first clade (1,1,1-TCA group) comprises Clones 1 and 2 and *ctrA, tmrA, cfrA* and *thmA* that have 2 or fewer total mismatches to the *cfrA* primer pair. The second clade (1,1-DCA group) comprises Clone 3 and *dcrA* and both are a perfect match to the *dcrA* primer pair. We thus refer to the genes amplified with the *cfrA* primers as “*cfrA*-like” genes, and those amplified with the *dcrA* primers as “*dcrA*-like” genes.

**Table 1.** Number of mismatches between dehalogenase gene sequences and primers.

| Target | *cfrA* forward | *cfrA* reverse | *dcrA* forward | *dcrA* reverse |
| --- | --- | --- | --- | --- |
| *Dhb* sp CF *cfrA* gene (JX282329.1) | 0 | 0 | 4 | 3 |
| *Desulfitobacterium* sp. PR *ctrA* gene (KF155162.1) | 2 | 0 | 4 | 3 |
| *Dhb* sp UNSWDHB *tmrA* gene (NZ_AUUR01000010 region: 19936..21303) | 0 | 0 | 4 | 3 |
| *Dhb* sp. SN *thmA* gene (KX344907.1) | 1 | 0 | 4 | 3 |
| Microcosm Clone1 | 1 | 0 | 4 | 4 |
| Microcosm Clone2 | 2 | 0 | 5 | 3 |
| *Dhb* sp DCA *dcrA* gene (JX282330.1) | 3 | 3 | 0 | 0 |
| Microcosm Clone3 | 3 | 3 | 0 | 0 |

Note: detailed alignments are shown in Fig. S7B.

### 3.3 Gene Abundances during Dechlorination in Groundwater Microcosms

We tracked the abundance of *Dehalobacter* (*Dhb*) 16S rRNA genes as well of the *cfrA*- and *dcrA*-like genes using qPCR in microcosm samples over time. Melt curves were examined, particularly for data close to or lower than the detection limit to confirm specific detection or not. When gene copy numbers were well above the detection limit, melt curves of samples and standards were identical (Supplementary Fig. S2).

The abundance the *cfrA*- and *dcrA-like* genes tracked dechlorination activity in a consistent and specific manner. As 1,1,1-TCA was dechlorinated to 1,1-DCA, the copy numbers of the *Dhb* 16S rRNA gene and the *cfrA*-like gene increased as well (Figs. 1 and S6). As 1,1-DCA was dechlorinated to CA in bottles #7, #9 and #18, the copy number of the *dcrA*-like gene increased. In bottles #8 (Fig. 1B) and #21 (Fig. S6C), when dechlorination was stuck at 1,1-DCA for more than 100 days, the copy number of the *dcrA*-like gene remained low (<10^7^ copies/ L). As soon as electron donor was added to these two bottles, 1,1-DCA was dechlorinated to CA and at the same time, the *dcrA*-like gene abundance increased by two orders of magnitude. Clearly, 1,1-DCA dechlorination was stalled due to lack of electron donor. We estimated the relative proportions of *Dehalobacter* carrying the *cfrA*-like and *dcrA*-like genes from the ratio of each functional gene to the 16S rRNA gene (%_*cfrA*-*Dhb*_ and %_*dcrA*-*Dhb*_). The %_*cfrA*-*Dhb*_ increased while 1,1,1-TCA was being converted to 1,1-DCA, and then declined while the %_*dcrA*-*Dhb*_ went up during the reduction of 1,1-DCA to CA (Fig. S8), confirming that these two genes clearly belonged to two different *Dhb* populations.

The coexistence of two highly similar but distinct *Dehalobacter* populations in the dechlorination of 1,1,1-TCA has only been reported in the study conducted by Tang *et al*. (Tang et al., 2012; Tang and Edwards, 2013B; Tang et al., 2016), where the authors speculated that the speciation into two highly specific strains in the ACT3 culture might be an artifact of laboratory cultivation over more than ten years in the presence of ample electron acceptor and donor. The current study suggests that this kind of speciation and selection can also take place *in situ* at a contaminated site. This realization also explains why dechlorination proceeded in a highly sequential manner where 1,1,1-TCA was first converted to 1,1-DCA before further dechlorination occurred. The second step required the growth of a distinct *Dhb* population which lagged behind and presumably competed with the first population for available donor. Consistently, the *dcrA*-containing population was at much lower abundance initially in all microcosms.

### 3.4 Biomass Yields Estimated from Microcosm Data

*Dehalobacter* cell yields, expressed as 16S rRNA copies per µmole chloride released for each dechlorination step, were estimated from microcosm data and compared to previously published values (Tables 2 and S6). Considering the many sources of possible errors in these measurements (including not accounting for cell decay, DNA extraction efficiency and inherent qPCR error), the results are reasonably consistent. Yields from microcosms with ZVI were slightly lower than those without ZVI, consistent with some abiotic transformation. Calculated cell yields from this study were similar to the yield reported for a pure culture of *Dehalobacter* sp. UNSWDHB (Wong et al., 2016), but are an order of magnitude lower than the yields reported for *Dehalobacter* sp TCA1 (Sun et al., 2002) and of *Dehalobacter* in the enrichment culture ACT3 (Grostern and Edwards, 2006B). The lower observed cell yields in this study are perhaps the result of incubating in groundwater only with no added vitamins.

**Table 2.**
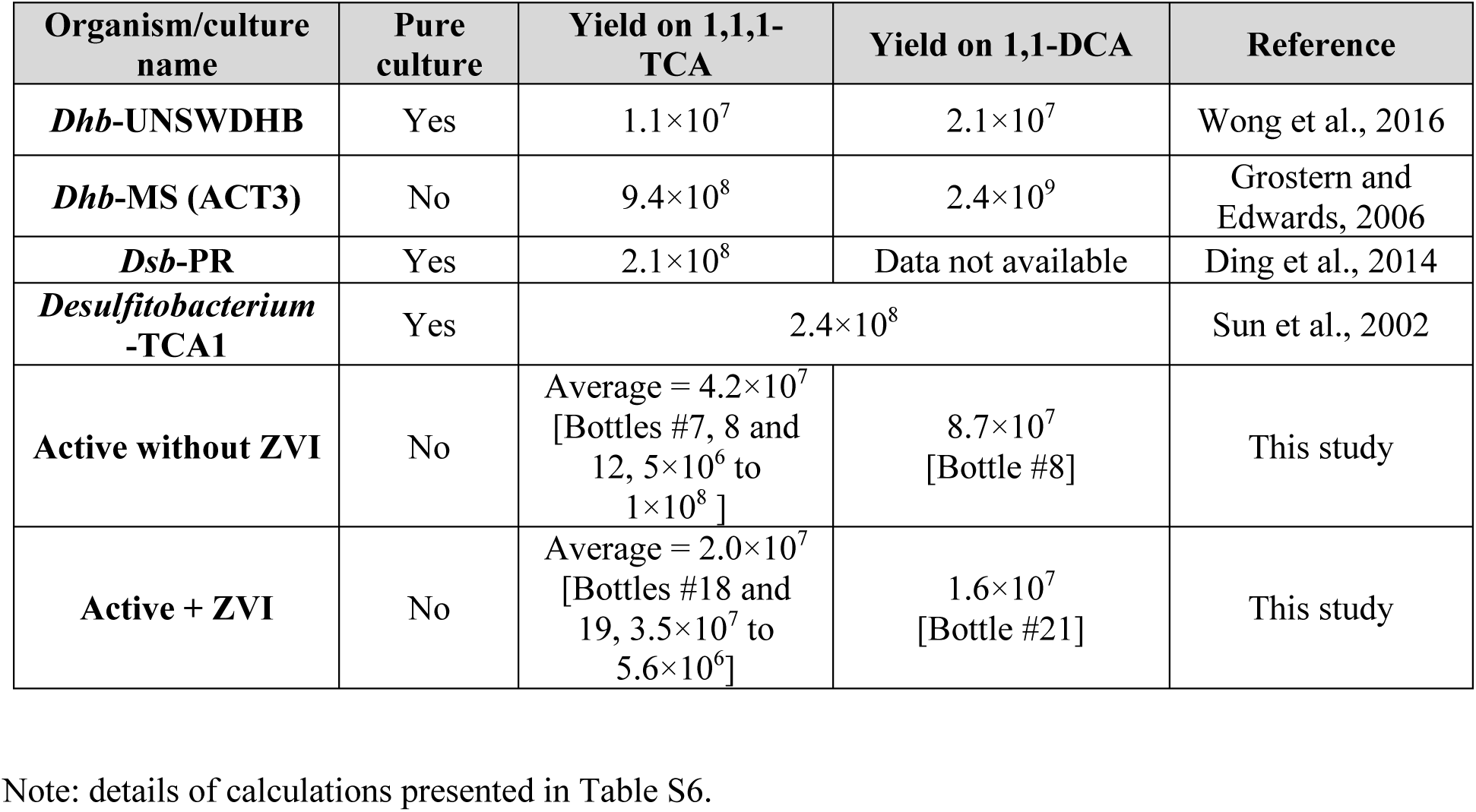
Summary of cell yields (gene copies / µmol Cl^−^ evolved) for organisms growing on 1,1,1-TCA and/or 1,1-DCA.

### 3.5 Analysis of Dechlorination Time Profiles and Gene Abundances in Field Samples

The abundances of *Dhb* 16S rRNA, *cfrA*-like and *dcrA*-like genes were determined in field samples to compare changes in gene abundances over time to changes in contaminant concentrations. Three wells were sampled extensively during this study, SW6, SW5 and MW202, all located in the plume area with high concentrations of 1,1,1-TCA (i.e., 0.45 – 1.2 mmol 1,1,1-TCA/L) prior to guar gum-ZVI injection. Because of the extent and variety of treatments applied to the site, including soil excavation, iron emplacement in trenches, injection and groundwater pumping, it was generally difficult to interpret the site concentration data.

SW6 was the only well with a clear pattern of transformation and a reasonable mole balance. After the guar gum-ZVI was injected to SW6, the decline of 1,1,1-TCA coincided with the generation of 1,1-DCA as well as increasing abundances of the *Dhb* 16S rRNA and *cfrA*-like genes (Fig. 2A). The abundance of the *dcrA*-like genes increased with a slight decline in the *cfrA*-like genes when 1,1-DCA was gradually converted to CA (after Day 250). These degradation profiles and correlations between the biomarker genes and the chlorinated ethane concentrations paralleled the observations from microcosm bottles. Assuming that the cell yield estimated from microcosms is applicable to the conditions in SW6, approximately 30% of the 1,1,1-TCA removed between Days 69 and 97 and 100% of that removed between Days 97 and 133 were attributed to dechlorination by *Dehalobacter* (Supplementary Table S7A). Towards the end of the field study, roughly 0.15 mmol/L CA was measured in groundwater from SW6, representing approximately 15% of the initial 1,1,1-TCA concentration. Since CA is strictly a biotic product, this 15% CA production provides a minimum estimate of the fraction dechlorinated by *Dehalobacter* carrying the *dcrA*-like gene. Therefore, biological dechlorination was responsible for a significant fraction of the observed dechlorination, from a minimum of 15% overall to a maximum of 100% at certain time intervals in SW6.

**Fig. 2.**
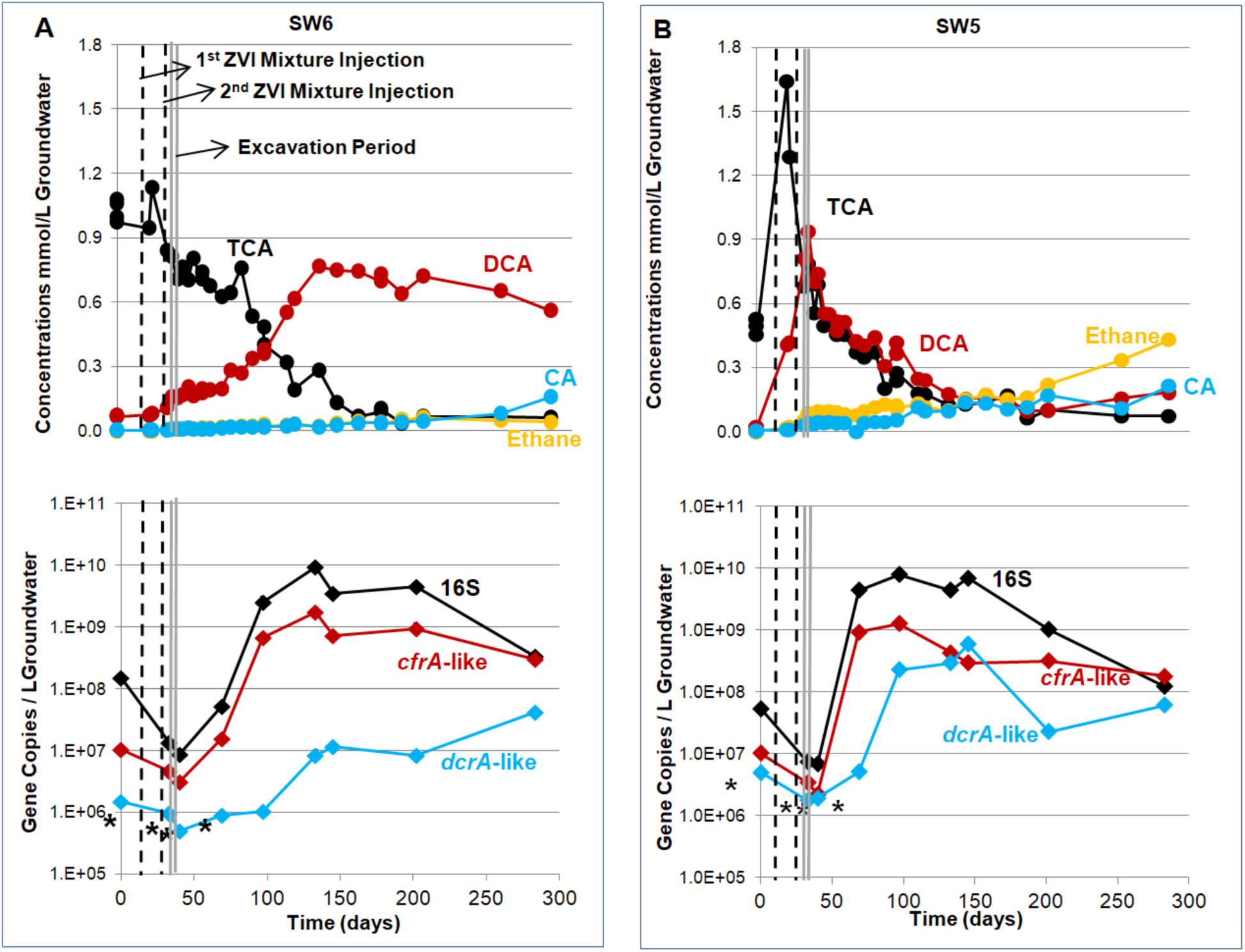
Chlorinated ethane concentrations and qPCR results in samples from wells SW6 (Panel A) and SW5 (Panel B). In each panel, the top figure shows chlorinated ethane concentrations (in circles, black for 1,1,1-TCA, red for 1,1-DCA, blue for CA, and yellow for ethane) and the bottom figure displays the qPCR results (in diamonds, black for the *Dhb* 16S rRNA gene, red for the *cfrA*-like gene, and blue for the *dcrA*-like gene). The values in the gene copies are the average of three technical replicates, and qPCR data points marked with **‘**\*’ are below the lowest detection limit or showed nonspecific amplification.

In monitoring wells SW5 (Fig. 2B) and MW202 (Supplementary Fig. S9), concentration data were difficult to interpret because the disappearance of 1,1,1-TCA did not coincide with stoichiometric production of 1,1-DCA. However, in SW5, the concentration of the *Dhb* 16S rRNA and *cfrA-like* genes increased by more than two orders of magnitude between Days 40 and 97 (Fig. 2B), revealing substantial biological dechlorination. Similar to the estimation done for SW6, based on the average cell yield in the microcosms and the qPCR results for SW5, biodechlorination accounted for roughly 60% of 1,1,1-TCA removed between days 40 and 97 in SW5 (Supplementary Table S7B). In MW202, biodechlorination was estimated to be about 10% of the 1,1,1-TCA removed between days 40 and 97 (Supplementary Table S7C). CA concentrations on Day 283 corresponded to 15% and 7% of the peak 1,1,1-TCA concentration in SW5 and MW202, respectively. Therefore, despite the noisy concentration data, the increase in the abundances of biomarker genes provided clear evidence that significant biological dechlorination was occurring in SW5 and MW202 during the periods monitored.

Our site data suggest that there needs to be at least ∼3×10^6^ copies of *Dehalobacter* 16S rRNA genes / L groundwater for measurable dechlorination of 1,1,1-TCA at the site. In addition, qPCR data from SW6 field samples (Fig. 2A) confirmed that indeed two distinct populations of *Dehalobacter* carrying two different RDase genes were present at the site, and each was responsible for one of the two dechlorinating steps. This observation is consistent with the observations in the groundwater microcosm studies presented earlier, and in the study of enrichment culture ACT-3 (Tang and Edwards, 2013B).

### 3.6 Microbial Communities in Microcosm and Site Samples

Amplicon sequencing based on the 16S rRNA gene was conducted to identify other organisms in the microcosm and field samples. These results were filtered to reveal organisms with max abundance > 5% in at least one sample (Table S8). *Dehalobacter* was certainly one of the dominant organisms in the microcosms (up to 34%). Other possible dechlorinating organisms include *Sulfurospirillum* (max 29%), *Dechloromonas* (max 7.6%) and *Geobacter* (max 3.6%, not shown in Table S8). Certain *Sulfurospirillum* and *Geobacter* species have been shown to dechlorinate chlorinated ethenes (Buttet et al., 2013; Sung et al., 2006), but not chlorinated ethanes. *Dechloromonas* was reported to dechlorinate chloroform (Lai et al., 2019). We examined the relationship between *Sulfurospirillum* abundance and 1,1,1-TCA and 1,1-DCA biodegradation. In microcosms (Figs S10A to S10C), *Sulfurosprillum* abundance did not increase as 1,1-DCA was dechlorinated to CA. In MW202 (Fig S10D), *Sulfurosprillum* abundance increased beyond Day 97 while no 1,1-DCA or CA was produced. Therefore, *Sulfurospirillum* was likely not responsible for the reduction of 1,1,1-TCA or 1,1-DCA in this study. Similarly, the trends in *Geobacter* and *Dechloromonas* did not correspond to the changes in the chlorinated ethane concentrations (Table S8). Thus, we can confidently state that *Dehalobacter* was responsible for biological 1,1,1-TCA and 1,1-DCA dechlorination in this study.

The amplicon sequencing data also provide some clues as to the likely organisms fermenting the guar gum polysaccharide, as well as an indication of the different selection pressures in the groundwater microcosm bottles versus the field samples. In the site samples, the relative abundance of several organisms increased substantially after guar gum injection (Tables S8). These included *Paludibacter, Trichococcus, Clostridium sensu stricto 1, Hypnocyclicus* and *Macellibacteroides*, all organisms known to metabolize carbohydrates to produce volatile fatty acids and/or simple alcohols (Murali et al., 2017; Wiegel et al., 2006; Roalkvam et al., 2015; Jabari et al., 2012; Liu et al., 2002). Interestingly, none of these organisms, except *Paludibacter*, became abundant in microcosm bottles. Rather, *Sulfurospirillum* increased substantially by day 49 in all batch bottles (Figs S10A to S10C) becoming one of the major organisms in the communities, yet as mentioned above, their abundance was not correlated to 1,1,1-TCA or 1,1-DCA dechlorination. Some *Sulfurospirillum* have been reported to ferment simple carbohydrates (Kruse et al., 2018; Kodama et al., 2007). Therefore, in our microcosms, *Sulfurospirillum* was the most likely fermenter to generate the H_2_ required by *Dehalobacter*. Moreover, it is clear that the lab microcosms were more reduced than the field samples, as shown by greater abundance of methanogens and other strict anaerobes.

### 3.7 Impact of Guar Gum and ZVI on Dechlorination of 1,1,1-TCA

Previous studies reported enhanced biodegradation of carbon tetrachloride, chloroform, 1,1,1-TCA and 1,2-dichloroethane after ZVI addition (Zemb et al., 2010; Zhou et al., 2014; Lee et al., 2015), while Xiu et al. (2010) found that the addition of nano-scale ZVI to a *Dehalococcoides*-containing methanogenic consortium initially inhibited trichloroethene dechlorination. In the current study, we did not observe adverse impacts of ZVI on *Dehalobacter* growth. Based on our parallel microcosm and site studies, the impact was most noticeable immediately after ZVI addition, with very rapid dechlorination of 1,1,1-TCA to 1,1-DCA. We also found that ZVI addition led to formation of some ethane and ethene from 1,1,1-TCA. While ethane concentration in MW202 and SW5 gradually increased to 0.2 – 0.4 mmol /L in over 280 days, ethane production in active bottles with ZVI addition mainly occurred on Day 1 and contributed to less than ∼10% of 1,1,1-TCA removal (Table S4B). Therefore, the reasons for the relatively high concentration of ethane observed in SW5 and MW202 are not well understood at this time. Overall, the main ZVI-related transformation production of 1,1,1-TCA was 1,1-DCA at the site, and further transformation of 1,1-DCA to CA was entirely biological.

Guar gum and its degradation products were a critical source of electron donor for the dechlorinating microbial communities in the field and in the microcosms. The monitoring well with lower COD concentration also contained lower abundance of *Dehalobacter* (e.g., MW202 between Day 50 and 100, Supplementary Fig. S11). In microcosms, the residual guar gum and fermentation products contained in the groundwater were sufficient to support biological dechlorination of 1,1,1-TCA and 1,1-DCA. Dechlorination stalled in only a few bottles, and in those cases, it was easily re-initiated by addition of electron donor.

## 4. Conclusions

This study established that injection of granular ZVI with guar gum supported a very active and sustained dechlorinating population that contributed to a large fraction of the observed dechlorination at the field site. Using an estimate of cell yield (copies *Dehalobacter* 16S rRNA genes per mol chloride released) in conjunction with qPCR abundance data we estimated the fraction of 1,1,1-TCA that was microbially dechlorinated versus that which was abiotically reduced by ZVI in actual field samples. By tracking the abundances of two specific functional genes (*cfrA*- and *dcrA*-like genes), we showed that two different *Dehalobacter* populations participated in the successive dechlorination reactions at the field site. One population was associated with dechlorination of 1,1,1-TCA to 1,1-DCA while the second was associated with dechlorination of 1,1-DCA to CA. Temporal and spatial trends in DNA-based biomarkers from groundwater samples provided an orthologous measure of biologically-mediated dechlorination reactions. Cell yields for obligate dechlorinating organisms such as *Dehalobacter* were used together with measured cell abundances to estimate contaminant transformation attributable to biodegradation. This is particularly valuable to deconvolute simultaneous abiotic and biotic transformation processes.

## Acknowledgements

We would like to acknowledge Pinchin Ltd. and NSERC for the financial support under the Industrial Postgraduate Scholarship program, the RENEW program and the INTEGRATE program funded by the Ontario Research Excellence Fund. We also thank Po-Hsiang Tommy Wang and Shuiquan Tang for offering the ACT3 culture and sharing their knowledge of the functional genes and primers.

## Supporting Information

Figures and tables in the Supplementary Information include detailed site information, microcosm setup, reaction recipe, and results of concentration data and sequence alignment that were not presented in the main text figures and tables.

## Abbreviations

1,1-DCA: 1,1-dichloroethane
1,1,1-TCA: 1,1,1-trichloroethane
CA: chloroethane
COD: chemical oxygen demand
*Dhb*: *Dehalobacter*
GC: gas chromatography
PCR: polymerase chain reaction
qPCR: quantitative polymerase chain reaction
RDases: reductive dehalogenases
ZVI: zero valent iron

## References

Bolyen, E., Rideout, J.R., Dillon, M.R., Bokulich, N.A., Abnet, C.C., Al-Ghalith, G.A., Alexander, H., Alm, E.J., Arumugam, M., Asnicar, F. et al., 2019. Reproducible, interactive, scalable and extensible microbiome data science using QIIME 2. Nat. Biotechnol. 37(8), 852–857.

Buttet, G.F., Holliger, C., Mailard, J., 2013. Functional genotyping of *Sulfurospirillum* spp. in mixed cultures allowed the identification of a new tetrachloroethene reductive dehalogenase. Appl Environ Microbiol. 79 (22), 6941–6947.

Cwiertny, D.M.; Bransfield, S.J.; Livi, K.J.; Fairbrother, D.H.; Roberts, A.L., 2006. Exploring the influence of granular iron additives on 1, 1, 1-trichloroethane reduction. Environ. Sci. Technol. 40, 6837–6843.

Damgaard, I.; Bjerg, P.L.; Bælum, J.; Scheutz, C.; Hunkeler, D.; Jacobsen, C.S.; Tuxen, N.; Broholm, M.M., 2013. Identification of chlorinated solvents degradation zones in clay till by high resolution chemical, microbial and compound specific isotope analysis. J. Contam. Hydrol. 146, 37–50.

Ding, C.; Zhao, S.; He, J., 2014. A *Desulfitobacterium* sp. strain PR reductively dechlorinates both 1,1,1-trichloroethane and chloroform. Environ. Microbiol. 16, 3387–3397.

Duchesneau, M. N.; Workman, R.; Baddour, F.R.; Dennis, P., 2007. Combined *Dehalobacter* and *Dehalococcoides* bioaugmentation for bioremediation of 1,1,1-trichloroethane and chlorinated ethenes. Proc. of the 9th international in situ and on-site bioremediation symposium. Baltimore, Maryland.

Fam. S.A.; Falatko, D.M.; McGillicuddy, G.; Pon, G.; Burkhardt, L.J.; Hone, J., 2012. Successful full-scale enhanced anaerobic dechlorination at a NAPL strength 1,1,1-TCA source area. Remed. J. 22, 33–47.

Fennelly, J.P.; Roberts, A.L., 1998. Reaction of 1, 1, 1-trichloroethane with zero-valent metals and bimetallic reductants. Environ. Sci. Technol. 32, 1980–1998.

Grostern, A.; Edwards, E.A., 2006. Growth of *Dehalobacter* and *Dehalococcoides* spp. during degradation of chlorinated ethanes. Appl. Environ. Microbiol. 72, 428–436.

Grostern, A.; Edwards, E.A. 2006B. A 1,1,1-trichloroethane-degrading anaerobic mixed microbial culture enhances biotransformation of mixtures of chlorinated ethenes and ethanes. Appl. Environ. Microbiol. 72, 7849–7856.

Gu, X.; Lu, S.; Li, L.; Qiu, Z.; Sui, Q.; Lin, K.; Luo, Q., 2011. Oxidation of 1,1,1-trichloroethane stimulated by thermally activated persulfate. Ind. Eng. Chem. Res. 50, 11029–11036.

Hug, L.A., Maphosa, F., Leys, D., Löffler, F.E., Smidt, H., Edwards, E.A., Adrian, L., 2013. Overview of organohalide-respiring bacteria and a proposal for a classification system for reductive dehalogenases. Philos. Trans. R. Soc. B. Biol. Sci. 368 (1616), 20120322.

Jabari, L., Gannoun, H., Cayol, J.L., Hedi, A., Sakamoto, M. Falsen, E., Ohkuma, M., Hamdi, M., Faugue, G. et al., 2012. *Macellibacteroides fermentans* gen. nov., sp. nov., a member of the family *Porphyromonadaceae* isolated from an upflow anaerobic filter treating abattoir wastewaters. Int. J. Syst. Evol. Microbiol. 62(10), 2522–2527.

Kocur, C.M.; Lomheim, L.; Boparai, H.K.; Chowdhury, A.I.; Weber, K.P.; Austrins, L.M.; Edwards, E.A.; Sleep, B.E.; O’Carroll, D.M., 2015. Contributions of abiotic and biotic dechlorination following carboxymethyl cellulose stabilized nanoscale zero valent iron injection. Environ. Sci. Technol. 49, 8648–8656.

Kodama, Y., Ha, L.T., and Watanabe, K., 2007. *Sulfurospirillum cavolei* sp. nov., a facultatively anaerobic sulfur reducing bacterium isolated from an underground crude oil storage cavity. Int. J. Syst. Evol. Microbiol. 57, 827–831.

Kruse, S., Goris, T., Westermann, M., Adrian, L., Diekert, G., 2018. Hydrogen production by *Sulfurospirillum* species enables syntrophic interactions of *Epsilonproteobacteria*. Nat. Commun. 9, 4872–2885.

Kruse, T., Maillard, J., Goodwin, L., Woyke, T., Teshima, H., Bruce, D., Detter, C., Tapia, R., Han, C., Huntemann, M., 2013. Complete genome sequence of *Dehalobacter restrictus* PER-K23 T. Stand. Genomic. Sci. 8, 375.

Lai, Y.S., Ontiveros-Valencia, A., Coskun, T., Zhou, C., Rittmann, B.E., 2019. Electron-acceptor loadings affect chloroform dechlorination in a hydrogen-based membrane biofilm reactor. Biotechnol Bioeng. 116, 1439–1448.

Liu, J.R., Tanner, R.S., Schumann, P., Weiss, N., McKenzie, C.A., Janssen, P.H., Seviour, E.M., Lawson, P.A., Allen, T.D., Seviour, R.J., 2002. Emended description of the genus *Trichococcus*, description of *Trichococcus collinsii* sp. nov., and reclassification of *Lactosphaera pasteurii* as *Trichococcus pasteurri* comb. Nov. and of *Ruminococcus palustris* as *Trichococcus palustris* comb. nov. in the low-G+C gram-positive bacteria. Int. J. Syst. Evol. Microbiol. 52, 1113–1126.

Lee, M.; Wells, E.; Wong, Y.K.; Koenig, J.; Adrian, L.; Richnow, H.H.; Manefield, M., 2015. Relative contributions of *Dehalobacter* and zero valent iron in the degradation of chlorinated methanes. Environ. Sci. Technol. 49, 4481–4489.

Lookman R, Bastiaens L, Borremans B, Maesen M, Gemoets J, Diels L., 2004. Batch-test study on the dechlorination of 1, 1, 1-trichloroethane in contaminated aquifer material by zero-valent iron. J. Contam. Hydrol. 74, 133–144.

Methods for the determination of inorganic substances in environmental samples; United States Environmental Protection Agency (EPA): EPA-600/R-93/100. August, 1993.

Molenda, O., Jacome, P., Cao, X., Nesbo, C.L.L., Tang, S.Q., Morson, N., Patron, J., Lomheim, L., Wishart, D.S.S. and Edwards, E.A., 2020. Insights into origins and function of the unexplored majority of the reductive dehalogenase gene family as a result of genome assembly and ortholog group classification. Environ. Sci.: Process. Impacts. 22(3), 663–678.

Murali, N., Srinivas, K., Ahring, B. K., 2017. Biochemical production and separation of carboxylic acids for biorefinery applications. Fermentation. 3, 22.

National Primary Drinking Water Regulations: 1,1,1-Trichloroethane; EPA-811-F-95-004r-T. United States Environmental Protection Agency (EPA). October, 1995.

Palau, J., Jamin, P., Badin, A., Vanhecke, N., Haerens, B., Brouyère, S., Hunkeler, D., 2016. Use of dual carbon–chlorine isotope analysis to assess the degradation pathways of 1,1,1-trichloroethane in groundwater. Water. Res. 92, 235–243.

Postiglione, J., Ferry, M., Quandt, L., Saappo, W., 2006. Bioremediation of TCE and TCA in groundwater by lactate injection. Proc. of the 5th international conference on remediation of chlorinated and recalcitrant compounds. Monterey, California.

Previdsa, M., 2016. In situ treatment of 1,1,1-trichloroethane by guar gum-stabilized zero valent iron injection: a field study. M.A.Sc. dissertation, University of Toronto, Toronto, Ontario.

Qiao, W., Puentes, L.A., Tang, X., Lomheim, L., Yang, M.I., Gaspard, S., Avanzi, I.R., Wu, J., Ye, S., Edwards, E.A., 2020. Microbial communities associated with sustained anaerobic reductive dechlorination of α-, β-, γ-, and δ-hexachlorocyclohexane isomers to monochlorobenzene and benzene. Environ. Sci. Technol. 54, 255–265.

Ritalahti, K.M., Hatt, J.K., Petrovskis, E., Löffler, F.E., 2009. Groundwater sampling for nucleic acid biomarker analysis. In Handbook of Hydrocarbon and Lipid Microbiology; Timmis, K. N. Eds.; Springer: Berlin, Germany. p3407–3418.

Roalkvam, I., Bredy, F., Baumberger, T., Pedersen, R-B., Steen, I.H., 2015. *Hypnocyclicus thermotrophus* gen. nov., sp. nov. isolated from a microbial mat in a hydrothermal vent field. Int. J. Syst. Evol. Microbiol. 65(12), 4521–4525.

Scheutz, C.; Durant, N.D.; Hansen, M.H.; Bjerg, P.L., 2011. Natural and enhanced anaerobic degradation of 1, 1, 1-trichloroethane and its degradation products in the subsurface–a critical review. Water. Res. 45, 2701–2723.

Sun, B.; Griffin, B.M.; Ayala-del-Rio, H.L.; Hashsham, S.A.; Tiedje, J.M., 2002. Microbial dehalorespiration with 1,1,1-trichloroethane. Science. 298, 1023–1025.

Sung, Y., Fletcher, K. E., Apkarian, R.P., Ramos-Hernández, N., Sanford, R.A., Mesbah, N.M., Löffler, F.E., 2006. *Geobacter lovleyi* sp. nov. strain SZ, a novel metal-reducing and tetrachloroethene-dechlorinating bacterium. Appl. Environ. Microbiol. 72(4), 2775–2782.

Tang, S., Edwards, E.A., 2013. Complete genome sequence of *Bacteroidales* strain CF from a chloroform-dechlorinating enrichment culture. Genome Announc. 1(6), e01066–13.

Tang, S.; Edwards, E.A., 2013B. Identification of *Dehalobacter* reductive dehalogenases that catalyse dechlorination of chloroform, 1,1,1-trichloroethane and 1,1-dichloroethane. Phil. Trans. R. Soc. B. 368:20120318.

Tang, S.; Gong, Y.; Edwards, E.A., 2012. Semi-automatic in silico gap closure enabled de novo assembly of two *Dehalobacter* genomes from metagenomic data. PLoS One, 7:e52038.

Tang, S.; Wang, P.H.; Higgins, S.A.; Löffler, F.E.; Edwards, E.A., 2016. Sister *Dehalobacter* genomes reveal specialization in organohalide respiration and recent strain differentiation likely driven by chlorinated substrates. Front. Microbiol. 7, 100.

Tobiszewski, M.; Namieśnik, J., 2012. Abiotic degradation of chlorinated ethanes and ethenes in water. Environ. Sci. Pollut. Res. 19, 1994–2006.

Toxicological profile for 1,1,1-trichloroethane. Agency for Toxic Substances and Disease Registry (ATSDR), U.S. Department of Health and Human Services, Public Health Service: Atlanta, GA, 2006.

Velimirovic, M.; Tosco, T.; Uyttebroek, M.; Luna, M.; Gastone, F.; De Boer, C.; Klaas, N.; Sapion, H.; Eisenmann, H.; Larsson, P.L., 2014. Field assessment of guar gum stabilized microscale zerovalent iron particles for in-situ remediation of 1,1,1-trichloroethane. J. Contam. Hydrol. 164, 88–99.

Wiegel, J., Tanner, R., Rainey, F.A., 2006. The Prokaryotes Volume 4: Bacteria: Firmicutes, *Cyanobacteria*. An Introduction to the family Clostridiaceae. M. Dworkin., S. Falkow., E. Rosenberg., K-H, Schleifer., E. Stackebrandt. Springer, U.S. P654–678.

Wong, Y.K.; Holland, S.I.; Ertan, H.; Manefield, M.; Lee, M., 2016. Isolation and characterization of *Dehalobacter* sp. strain UNSWDHB capable of chloroform and chlorinated ethane respiration. Environ. Microbiol. 18, 3092–3105.

Wu, X.; Lu, S.; Qiu, Z.; Sui, Q.; Lin, K.; Du, X.; Luo, Q., 2014. The reductive degradation of 1, 1, 1-trichloroethane by Fe (0) in a soil slurry system. Environ. Sci. Pollut. Res. 21, 1401–1410.

Xiu, Z.; Jin, Z.; Li, T.; Mahendra, S.; Lowry, G.V.; Alvarez, P.J., 2010. Effects of nano-scale zero-valent iron particles on a mixed culture dechlorinating trichloroethylene. Bioresour. Technol. 101, 1141–1146.

Zemb, O.; Lee, M.; Low, A.; Manefield, M., 2010. Reactive iron barriers: A niche enabling microbial dehalorespiration of 1, 2-dichloroethane. Appl. Microbiol. Biotechnol. 88, 319–325.

Zhao, S.; Rogers, M. J; He, J., 2017. Microbial reductive dehalogenation of trihalomethanes by a *Dehalobacter*-containing co-culture. Appl. Microbiol. Biotechnol. 101, 5481–5492.

Zhou, Y.; Yang, J.; Wang, X.; Pan, Y.; Li, H.; Zhou, D.; Liu, Y.; Wang, P.; Gu, J.; Lu, Q., 2014. Bio-beads with immobilized anaerobic bacteria, zero-valent iron, and active carbon for the removal of trichloroethane from groundwater. Environ. Sci. Pollut. Res. 21, 11500–11509.

