## Supplemental information for "Using cell yields and qPCR to estimate biotic contribution to 1,1,1-trichloroethane dechlorination at a field site treated with granular zero valent iron and guar gum"

**Table S1. Historical VOC data from wells MW202, SW5 and SW6**

| Unit: µg/L | SW5 | | | | SW6 | | | | |
|---|---|---|---|---|---|---|---|---|---|
| Date | 28-Aug-99 | 1-Jul-04 | 10-Jul-11 | 28-Jul-12 | 28-Aug-99 | 1-Jul-04 | 10-Jun-11 | 28-Jun-12 | 25-Jun-14 |
| Acetone | NA | 6200 | <50000 | <100000 | NA | 5800 | <50000 | <50000 | <100000 |
| 1,1-Dichloroethane | 2900 | 1600 | 3300 | 5800 | 2200 | 140 | 2500 | 1300 | 3200 |
| 1,1-Dichloroethene | 2000 | 1600 | 3300 | 5800 | 2400 | 920 | 5800 | 6800 | 7600 |
| Tetrachloroethene | 50 | 44 | <500 | <2000 | 100 | 79 | <500 | <500 | <1000 |
| 1,1,1-Trichloroethane | 50000 | 33000 | 90000 | 150000 | 61000 | 12000 | 95000 | 100000 | 160000 |
| Trichloroethene | 160 | 55 | <500 | <1000 | 200 | 92 | <500 | <500 | <1000 |

**MW202**

| Unit: µg/L | 7-Jan-04 | 30-Oct-07 | 02-Feb-08 | 1-Ar-09 | 15-Jun-10 | 10-Jun-11 | 27-Jun-12 | 25-Jun-14 |
|---|---|---|---|---|---|---|---|---|
| Tetrachloromethane | 78.8 | <8 | 160 | <160 | <2000 | <20 | 3.9 | <5000 |
| Chloroethane | <20 | 51 | 100 | | <160 | <4000 | NA | NA |
| Trichloromethane | 84.8 | 26 | 70 | <160 | <2000 | <20 | 16 | <5000 |
| 1,1-Dichloroethane | 42300 | 6600 | 26000 | 110000 | 35000 | 1800 | 8000 | 110000 |
| 1,2-Dichloroethane | 527 | 200 | 310 | 2100 | <4000 | 22 | 100 | <10000 |
| 1,1-Dichloroethene | 7650 | 4500 | 25000 | 160000 | 4400 | 570 | 2200 | 12000 |
| Cis-1,2-Dichloroethene | 21 | 63 | 83 | <160 | <2000 | 11 | 34 | <5000 |
| Trans-1,2-Dichloroethene | <10 | <8 | 51 | <160 | <2000 | <0.5 | 1.1 | <5000 |
| Dichloromethane | 60500 | 20000 | 23000 | 270000 | 64000 | 1900 | 12000 | 160000 |
| Tetrachloroethene | 81.6 | 31 | 150 | <160 | <2000 | 8.8 | 22 | <5000 |
| Toluene | 57.6 | 21 | 130 | <160 | <4000 | <20 | <100 | <10000 |
| 1,1,1-Trichloroethane | 714000 | 130000 | 2600000 | 1500000 | 540000 | 28000 | 110000 | 930000 |
| 1,1,2-Trichloroethane | 44.8 | 18 | <20 | <160 | <4000 | 2.4 | 7.3 | <10000 |
| Trichloroethene | 4240 | 1400 | 5000 | 6100 | 2700 | 150 | 640 | <5000 |
| Vinyl Chloride | <20 | <6.8 | 26 | <136 | <4000 | <0.2 | 1.6 | <10000 |

**Table S2. Details of guar gum-ZVI injections**

| | |
|---|---|
| **Granular ZVI information** | |
| **Cast iron aggregate size** | 8/50, 0.37 - 2.36mm |
| **ZVI manufacturer** | Peerless Metal Powders & Abrasive |
| **Guar gum matrix information** | |
| **Matrix components** | Water, guar gum gel (Biofrac G), cross-linker (Biofrac X) and a caustic pH buffer |
| **Guar gum matrix manufacturer** | Frac Rite Environmental Ltd, Alberta, Canada |
| **Drilling information** | |
| **Drilling method** | Direct push drilling with Geoprobe Model 6620DT |
| **Fracturing pressure** | 600 - 900 psi |
| **Propagation pressure** | 150 psi |
| **Injection in the plume area** | |
| **Injection intervals per injection point** | 0.75 m intervals, between 2.5 and 8.5 meters below ground surface (mbgs) |
| **Injection point spacing** | 7 m |
| **Total ZVI injected to plume area** | 17000 kg |
| **ZVI injected per injection interval** | 226 kg |
| **Total mixture injected** | 30400 L |
| **Mixture injected per interval** | 400 L |
| **Additional guar gum injected to propagate the fractures** | 5600 L |
| **Soil excavation** | |
| **Number of trenches** | 3 |
| **Width of trench** | 1 m |
| **Average trench depth** | 6.5 m |
| **Total impacted soil removed** | 410 tonnes |
| **Backfilled ZVI to trenches** | 20000 kg |
| **Depth of ZVI in backfill** | 2 - 6.5 mbgs |

Note: a series of confirmatory boreholes were drilled between injection points to ensure a sufficiently uniform ZVI distribution. Venting of gel and fine iron particulates to the surface was noted up to 10 meters directly upgradient of multiple injection points at fracture depths as low as 8.5 mbgs.

**Table S3. Groundwater microcosm setup**

| Description | Bottle #[1] | Groundwater[2] added as inoculum and $e^-$ donor (mL) | ACT3 culture added (mL) | Neat 1,1,1-TCA added (µL) | ZVI added (g) | Autoclaved |
|---|---|---|---|---|---|---|
| Sterile | 1, 2, 3 | 150 | 0 | 4.25 | 0 | Yes |
| Sterile + ZVI[3,4] | 4, 5, 6 | 150 | 0 | 4.25 | 1 | Yes |
| Active without ZVI | 7, 8[6], 9, 10, 11, 12, 13 | 150 | 0 | 4.25 | 0 | No |
| Active + ZVI | 17, 18, 19, 20, 21[6], 22 | 150 | 0 | 4.25 | 1 | No |
| ACT3 + ZVI | 14, 15, 16 | 150 | 0.1 | 4.25 | 1 | No |
| Active without TCA and ZVI[5] | 23 | 150 | 0 | 0 | 0 | No |

Notes:

1. The bottles listed here includes the entire list of bottles set up in this groundwater microcosm experiment. Greyed bottle numbers are for results not reported here. Bottle #6 leaked and bottle #15 cracked, so they are not presented. Other bottles highlighted in grey, including bottles #10, #11, #13, #17, #20 and #22 behaved very similarly to other bottles in the same set, therefore we do not show this data to simplify presentation of results.
2. Groundwater collected on day 202 from well SW6 at the site served as the source of inoculum, nutrients and carbon (i.e., from residual guar gum and degradation products).
3. After the addition of groundwater and ZVI, the Sterile + ZVI bottles were placed in autoclave with aluminum foil wrapping the bottle opening. The bottle caps were disinfected separately by wiping with ethanol. After autoclaving at 120 °C and 18 psi for 40 minutes, these hot bottles were capped immediately with the disinfected screw caps and purged with hot $N_2/CO_2$ gas mix filtered with sterile 0.22 µm filters (Millipore, Billerica, MA) for 30 minutes to re-establish anaerobic conditions prior to adding 1,1,1-TCA. The headspace pressure was 1atm at room temp. prior to 1,1,1-TCA addition.
4. The outer surface of the granular ZVI in the Sterile + ZVI bottles might be partially oxidized during autoclaving, as the bottles were exposed to open atmosphere. However, since the solubility of oxygen in water was relatively small at high temperature, we believed that the reactivity of the ZVI did not change substantially. Indeed, we also used alternative approaches to sterilize another two sets of 'Sterile + ZVI' controls, one using gamma irradiation and the other using sodium azide and mercuric chloride. These bottles were always kept anaerobically during and after assay setup. All three sets of 'Sterile+ ZVI' controls, regardless of the different means of sterilization, showed similar trends: 1,1,1-TCA was completely reduced to 1,1-DCA as the final and dominant daughter product; reduction took place immediately after assay setup; and the total moles of C2 compounds decreased initially.
5. The single 'Active without TCA and ZVI' microcosm was set up in a serum bottle with rubber stopper and metal crimp instead of a glass bottle with the Mininert cap, and neither 1,1,1-TCA nor ZVI was added to this bottle.
6. In bottles #8 and #21, on Day 140 when 1,1-DCA concentration had not changed for more than 90 days, ethanol and lactate were added to these two bottles. The addition of ethanol and lactate was at five times excess as compared to the remaining 1,1-DCA on the electron equivalent basis in order to assess if donor was limiting dechlorination.

**Table S4. Methane, ethane and ethene production in microcosms**

**Table S4A. Methane production in each bottle on Day 245**

| Case | Bottle # | mmol of methane on Day 245 | % of maximum theoretical methane production |
|---|---|---|---|
| Sterile | 1 | 0 | 0 |
| | 2 | 0 | 0 |
| | 3 | 0 | 0 |
| Sterile + ZVI | 4 | 0.004 | 0.2 |
| | 5 | 0.003 | 0.2 |
| Active without ZVI | 7 | 0.963 | 47.4 |
| | 8 | 0.056 | 2.8 |
| | 9 | 1.123 | 55.3 |
| | 12 | 0.001 | 0.1 |
| Active + ZVI | 18 | 0.004 | 0.2 |
| | 19 | 0.006 | 0.3 |
| | 21 | 0.012 | 0.6 |
| ACT3 + ZVI | 14 | 0.368 | 18.1 |
| | 16 | 0.477 | 23.5 |
| Active without ZVI or 1,1,1-TCA | 23 | 1.844 | 90.8 |

**Table S4B. Ethane and ethene production in each microcosm bottle on Day 1 and Day 245**

| Case | Bottle # | mmol of ethane | | mmol of ethene | |
|---|---|---|---|---|---|
| | | Day 1 | Day 245 | Day 1 | Day 245 |
| Sterile | 1 | 0 | 0 | 0 | 0 |
| | 2 | 0 | 0 | 0 | 0 |
| | 3 | 0 | 0 | 0 | 0 |
| Sterile + ZVI | 4 | 0.005 | 0.006 | 0.001 | 0.003 |
| | 5 | 0.004 | 0.005 | 0.001 | 0.003 |
| Active without ZVI | 7 | 0 | 0 | 0 | 0 |
| | 8 | 0 | 0 | 0 | 0 |
| | 9 | 0 | 0 | 0 | 0 |
| | 12 | 0 | 0 | 0 | 0 |
| Active + ZVI | 18 | 0.001 | 0.003 | 0.001 | 0.002 |
| | 19 | 0.002 | 0.004 | 0.001 | 0.002 |
| | 21 | 0.005 | 0.003 | 0.002 | 0.002 |
| ACT3 + ZVI | 14 | 0.003 | 0.002 | 0.001 | 0.001 |
| | 16 | 0.003 | 0.002 | 0.001 | 0.001 |
| Active without ZVI or 1,1,1-TCA | 23 | 0 | 0 | 0 | 0 |

Note that ethene and ethane were only detected in bottles with ZVI

**Table S5. Percent similarity for reductive dehalogenase sequences**

Shown are amino acid similarity (top, yellow, length = 411 amino acids) and nucleotide similarity (bottom, green, length = 1236 nucleotides) between reductive dehalogenase genes (from known dechlorinating organisms) and clone sequences in this study.

Red numbers indicate the highest similarity scores of our clone sequences as compared to the reference genes.

| Target | ***Dhb* sp CF *cfrA* Gene** | ***Dsf* sp. PR *ctrA* Gene** | ***Dhb* sp UNSWDHB *tmrA* Gene** | ***Dhb* sp. THM1 *thmA* Gene** | **Microcosm Clone 1** | **Microcosm Clone 2** | ***Dhb* sp DCA *dcrA* Gene** | **Microcosm Clone 3** |
|---|---|---|---|---|---|---|---|---|
| ***Dhb* sp CF *cfrA* Gene** | | 94.2 | 95.9 | 95.4 | 95.1 | 93.9 | 94.9 | 94.6 |
| ***Dsf* sp. PR *ctrA* Gene** | 97 | | 94.4 | 98.1 | 97.1 | 94.4 | 93.7 | 93.4 |
| ***Dhb* sp UNSWDHB *tmrA* Gene** | 98.2 | 97.4 | | 95.9 | 95.4 | 94.2 | 95.1 | 94.9 |
| ***Dhb* sp. THM1 *thmA* Gene** | 97.6 | 99.4 | 98.1 | | **98.3** | **96.1** | 95.1 | 94.9 |
| **Microcosm Clone 1** | 97.5 | 98.9 | 97.7 | **99.4** | | 95.1 | 93.9 | 93.7 |
| **Microcosm Clone 2** | 97.2 | 97.6 | 97.5 | **98.1** | 97.6 | | 93.7 | 93.4 |
| ***Dhb* sp DCA *dcrA* Gene** | 97.7 | 96.8 | 97.7 | 97.5 | 97 | 96.9 | | **99.5** |
| **Microcosm Clone 3** | 97.7 | 96.8 | 97.7 | 97.5 | 97 | 96.9 | **99.8** | |

.

**These three larger Tables are provided in a separate Excel file:**

Table S6. Detailed calculation of cell yields

Table S7. Estimation of %TCA biodegraded at the site using yields and qPCR data

Table S8. Major organism features found in amplicon (Illumina) sequencing

**Supporting Text S1: Site stratigraphy**

The site stratigraphy consisted of reworked silty clay till containing lenses of silt and fine sand from zero to three meters below ground surface (mbgs), native clay till from approximately 3 to 17 mbgs and an underlying deep sand aquifer starting at 17 mbgs. Chlorinated compounds were present in a perched groundwater flow system in the native clay till, but not in the deep water table in the underlying sand aquifer. The hydraulic conductivity of the overburden unit ranged from approximately $10^{-7}$ to $10^{-8}$ m/sec, and $10^{-6}$ m/s in the more permeable lenses (Previdsa, 2016). A pump and treat system was operated between 1995 and 2015, and 4800 tonnes of impacted soil were excavated around year 2000.

**Supporting Text S2. qPCR recipe and programs**

Calibration curves were constructed using known concentrations of plasmid DNA containing the corresponding 16S rRNA gene or the reductive dehalogenase gene insert. Each calibration curve consisted of serial ten-fold dilutions of the plasmid DNA for the gene of interest, from approximately $10^8$ to $10^2$ copies /µL.

DNA samples (2 µL, diluted field/microcosm/standard plasmid) were mixed with 18 µl of mastermix that included the following: 10 µL 10× SsoFastTM EvaGreen® Supermix (BioRad, California, US, #1725202), 0.5 µL 10mM forward primer, 0.5 µL 10mM reverse primer, 7 µL UltraPure DNase/Rnase-free distilled water (Life Technology, California, US, #10977015). All primers were purchased from Sigma-Aldrich (Missouri, US). The reaction was run in a BioRad CFX96 Touch™ Real-Time PCR Detection System with the following program settings: initial denaturation for 2 min at 98°C; 40 cycles of denaturation at 98°C for 5 sec, annealing at 62.5°C (for the *Dhb* 16S rRNA primer) or 60°C (for the *cfrA* and *dcrA* primer sets) for 10 sec; a final melting curve analysis from 65 to 95°C, measuring fluorescence every 0.5°C. The analysis was performed using the Bio-Rad CFX Manager 3.1 program. The amplification efficiency values were 88%, 89.9% and 95.8% for *Dhb* 16S rRNA gene, *cfrA* gene and *dcrA* gene respectively. The lowest quantification limits in our runs were 370, 125 and 40 copies/ reaction (i.e., per µL DNA template added) for *Dhb* 16S rRNA gene, *cfrA* gene and *dcrA* gene correspondingly. The melt curves are presented in Fig S2.

**Supporting Text S3: Amplicon sequencing of the site samples and microcosm samples**

The DNA extracted from site samples and groundwater microcosm samples were sent to McGill University and Genome Quebec Innovation Center for 16S rRNA gene amplicon sequencing using the Illumina MiSeq PE300 platform with the modified versions of primers 926f and 1392r (926f-modified: AAACTYAAAKGAATWGRCGG; 1392r-modified: ACGGGCGGTGWGTRC) (Qiao et al., 2020). The sequencing data were processed using the QIIME2 pipeline (Bolyen et al., 2019). After the raw sequences were imported to QIIME2 as an artifact, forward and reversed primers were removed using the 'qiime cutadapt trim-paired' command. Then the trimmed sequences were filtered, denoised, merged (i.e., joining forward and reverse reads), chimera-checked and clustered into different features based on amplicon sequence variance (ASV) using the Dada2 package with the 'qiime dada2 denoise-paired' command, with the following settings: p-trunc-len-f = 260, p-trunc-len-r = 230. p-max-ee = 2. The representative sequences of the features were then assigned with taxonomy using the 'feature-classifier classify-sklearn' command with the Silva-132-99-nb classifier trained to match our 926f and 1392r primer set. The final biom table along with the taxonomy assignment was exported as a tsv file and the percentage of each feature and genus was calculated in Microsoft Excel.

The following sample accession numbers were obtained for the raw amplicon sequencing reads submitted to the NCBI SRA: SAMN14988330 for MW202_Day0, SAMN14988331 for MW202_Day145, SAMN14988332 for MW202_Day202, SAMN14988333 for MW202_Day33, SAMN14988334 for MW202_Day40, SAMN14988335 for MW202_Day69, SAMN14988336 for MW202_Day97, SAMN14988337 for NoZVI_B12_Day139, SAMN14988338 for NoZVI_B12_Day181, SAMN14988339 for NoZVI_B12_Day49, SAMN14988340 for NoZVI_B7_Day139, SAMN14988341 for NoZVI_B7_Day49, SAMN14988342 for NoZVI_B8_Day139, SAMN14988343 for NoZVI_B8_Day181,

SAMN14988344 for NoZVI_B8_Day49, SAMN14988345 for NoZVI_B9_Day49, SAMN14988346 for SW5_Day0, SAMN14988347 for SW5_Day133, SAMN14988348 for SW5_Day145, SAMN14988349 for SW5_Day202, SAMN14988350 for SW5_Day33, SAMN14988351 for SW5_Day40, SAMN14988352 for SW5_Day69, SAMN14988353 for SW5_Day97, SAMN14988354 for SW6_Day0, SAMN14988355 for SW6_Day133, SAMN14988356 for SW6_Day145, SAMN14988357 for SW6_Day202, SAMN14988358 for SW6_Day33, SAMN14988359 for SW6_Day40, SAMN14988360 for SW6_Day69, SAMN14988361 for SW6_Day97, SAMN14988362 for ZVI_B18_Day139, SAMN14988363 for ZVI_B18_Day49, SAMN14988364 for ZVI_B19_Day139, SAMN14988365 for ZVI_B19_Day181, SAMN14988366 for ZVI_B19_Day49, SAMN14988367 for ZVI_B21_Day139, SAMN14988368 for ZVI_B21_Day181, and SAMN14988369 for ZVI_B21_Day49, where 'ZVI' stands for the set of 'Active + ZVI' and 'NoZVI' stands for the set of 'Active without ZVI'.

**Supporting Text S4: *rdhA* clone sequences**

Red: forward primer binding site (outside of coding region)
Green: *rdhA* coding region (incomplete at 3' end)
Purple: *rdhA* reverse primer binding site

>Rdases_Gene_Clone1 [*Dehalobacter*] [from groundwater_microcosm] *cfrA* gene-like, partial gene (5' to 3')
AAGAGATTGTAGAAGCAGCGGCTGCTCTGATGCACTCGAAAGGATATGAAAACACCAAACT
CAGCGATATTCTAAAGGAGGGTCAACTTATGGAAAAAGAAAAATGTAACAATGATGAGCCG
GCAACAATGGACAAGGAAAAAAGTAACAACGATAAGCCGGCAACAAAAATTAATCGCAGAC
GATTCCTTAAATTTGGAGCTGGAGCTTCTTCAGGTATTGCAATTGCCGCTGCAGCTACTGCAT
TGGGAGGGAAATCACTTATCGATCCCAAACAGGTATATGCTGGAACGGTCAAGGAACTGGAT
GAACTTCCCTTTAATATCCCGGCAGACTACAAACCGTTTACCAATCAAAGGAATATATTTGGC
CAGGCTGTATTGGGAGTACCAGAACCTCTAGCACTTGTAGAGCGTTTTGATGAAGTAAGATG
GAATGGTTGGCAGACAGATGGTTCGCCCGGTCTTACTGTACTTGATGGTGCGGCTGCTCGTGC
AAGCTTTGCCGTCGATTATTATTTTAACGGGGAAAATAGCGCCTGCAGGGCCAATAAAGGTT
TTTTTGAATGGCATCCCAAAGTGCCCGAGCTGAACTTTAGGTGGGGCGATCCGGAGAGAAAT
ATTCATTCCCCCGGTGTAAAAAGTGCCGAAGAAGGAACGATGGCAGTAAAAAGAATGGCTA
GATTTTTCGGCGCTGCTAAAGCTGGGATAGCGCCTTTTGACAAACGTTGGGTTTTTACTGAAA
CGGCTGCCTTTGTTAAAACGCCTGAGGGTGAAAGTCTGAAATTTATCCCTCCGGATTTTGGGT
TTGAGCCCAAGCACGTAATATCGATGATTATCCCACAGTCGCTAGAAGGAGTAAAGAGTGCC
CCGTCCTTTTTAGGATCGTCTGAATATGGATTAAGTTGTGCCCAGTATGGATATGCTCCATTC
GGTTTATCCATGTTTATTAAAGATCTGGGATATCATGCGGTTCCAATCGGAGCTGACAGTGCA
TTAGCTATACCTATAGCTATTCAGGCGGGTCTGGGGGAATACAGCAGGTCGGGGCTAATGAT
TACGCCTGAATTTGGTTCAAATTTTAGACTCTGTGAAGTATTTACTGACATGCCTTTAAATCA
TGATAAACCTATTTCATTCGGAGTAACTGAATTTTGCAAAACCTGCAAAAAATGCGCTGAAG

AATGCGTCCCTCAAGCTATTAGCTATGAAGATCCTACCATTGATGGACCTCGTGGGCAAATG
CAAAATTCGGGAGTAAAGAGATGGTATGTTGACCCGGTGAAGTGCCTTGAATTCTGGTCGCG
TGATAACGTCAGAAACTGCTGCGGAGCTTGTATAGCTGCTTGCCCATTTACTAAG

Translation of the coding region of Clone1:
MDKEKSNNDKPATKINRRRFLKFGAGASSGIAIAAAATALGGKSLIDPKQVYAGTVKELDELPFN
IPADYKPFTNQRNIFGQAVLGVPEPLALVERFDEVRWNGWQTDGSPGLTVLDGAAARASFAVDY
YFNGENSACRANKGFFEWHPKVPELNFRWGDPERNIHSPGVKSAEEGTMAVKRMARFFGAAKA
GIAPFDKRWVFTETAAFVKTPEGESLKFIPPDFGFEPKHVISMIIPQSLEGVKSAPSFLGSSEYGLSC
AQYGYAPFGLSMFIKDLGYHAVPIGADSALAIPIAIQAGLGEYSRSGLMITPEFGSNFRLCEVFTD
MPLNHDKPISFGVTEFCKTCKKCAEECVPQAISYEDPTIDGPRGQMQNSGVKRWYVDPVKCLEF
WSRDNVRNCCGACIAACPFTK

>Rdases_Gene_Clone2 [*Dehalobacter*] [from groundwater_microcosm] *cfrA* gene-like, partial gene (5' to 3')
AAGAGATTGTAGAAGCAGCGGCTGCTCTGATGCACTCGAAAGGATATGAAAACACCAAACT
CAGCGATATTCTAAAGGAGGGTCAACTTATGGAAAAAGAAAAATGTAACAATGATGAGCCG
GCAACAATGGACAAGGAAAAAAGTAACAACGATAAGCCGGCAACAAAAATTAATCGCAGAC
GATTCCTTAAATTTGGAGCTGGAGCTTCTTCAGGTATTGCAATTGCCGCTGCAACTACTGCAT
TGGGAGGGAAATCACTTATCGATCCCAAACAGGCAAATGCTGGAACGGTCAAGGAACTGGA
TGAACTTCCCTTTGATATCCCGGCAGACTACAAACCGTTTACCAATCAAAGGAATATATATG
GCCAAGCTATATTGGGAGTACCAGAACCTCTAGCACTTGAAGAGCGTTTTAATGAAGTAAGA
TGGAATGGTTGGCAGACAGATGGTTCGCCCGGTCTTACTGTACTTGATGGTGCGGCTGCTCGT
GCAAGCTTTGCCGTTGATTATTATTTTAACGGGGAAAATAGCGCCTGCAGGGCCAATAAAGG
TTTTTTTGAATGGCATCCCAAAGTGCCCGAGCTGAACTTTAGGTGGGGCGATCCGGAGAGAA
ATATTCATTCCCCCGGTGTAAAAAGTGCCGAAGAAGGAACGATGGCAGTAAAAAGAATGGC
TAGATTTTTCGGCGCTGCTAAAGCTGGGATAGCGCCTTTTGACAAACGTTGGGTTTTTACTGA
AACGGCTGCCTTTGTTAAAACGCCTGAGGGTGAAAGTCTGAAATTTATCCCTCCGGATTTTGG
GTTTGAGCCCAAGCACGTAATATCGATGATTATCCCACAGTCGCTAGAAGGAGTAAAGTCTG
CCCCGTCCTTTTTAGGATCAGTTGAATATGGATTAAGTTTTGCCCAGTTTGGATATGCTGCAT
TCGGTTTATCCATGTTTATTAAAGATCTGGGATATCATGCGGTTCCAATCGGAGCTGACAGTG
CATTAGCTATACCTATAGCTATTCAGGCGGGTCTGGGGGAATACAGCAGGTCGGGGCTAATG
ATTACGCCTGAATTTGGTCCAAATGTTAAACTCTGTGAAGTATTTACTGACATGCCTTTAAAT
CATGATAAACCTATTTCATTCGGAGTAACTGAATTTTGCAAAACCTGCAAAAAATGCGCTGA
AGCATGCGCCCCTCAAGCTATTAGCTATGAAGATCCTACCATTGATGGACCTCGTGGGCAAA
TGCAAAATTCGGGAATAAAGAGATGGTATGTTGACCCGGTGAAGTGCTTGGAATTCTTTTCG
CGTGATAACGTCAAAAACTGCTGCGGAGCTTGTATAGCTGCTTGCCCATTTACTAAG

Translation of the coding region of Clone2:
MDKEKSNNDKPATKINRRRFLKFGAGASSGIAIAAATTALGGKSLIDPKQANAGTVKELDELPFDI
PADYKPFTNQRNIYGQAILGVPEPLALEERFNEVRWNGWQTDGSPGLTVLDGAAARASFAVDYY
FNGENSACRANKGFFEWHPKVPELNFRWGDPERNIHSPGVKSAEEGTMAVKRMARFFGAAKAG
IAPFDKRWVFTETAAFVKTPEGESLKFIPPDFGFEPKHVISMIIPQSLEGVKSAPSFLGSVEYGLSFA
QFGYAAFGLSMFIKDLGYHAVPIGADSALAIPIAIQAGLGEYSRSGLMITPEFGPNVKLCEVFTDM
PLNHDKPISFGVTEFCKTCKKCAEACAPQAISYEDPTIDGPRGQMQNSGIKRWYVDPVKCLEFFSR
DNVKNCCGACIAACPFT

>Rdases_Gene_Clone3 [*Dehalobacter*] [from groundwater_microcosm] *dcrA* gene-like, partial gene (5’ to 3’)

AAGAGATTGTAGAAGCAGCGGCTGCTCTGATGCACTCGAAAGGACATGAAAACACCAAACT
CAGCGATATTCTAAAGGAGGGTCAACTTATGGAAAAAGAAAAATGTAACAATGATGAGCCG
GCAACAATGGACAAGGAAAAAAGTAACAACGATAAGCCGGCAACAAAAATTAATCGCAGAC
AATTCCTTAAATTTGGAGCTGGAGCTTCTTCGGGTATTGCAATTGCCACTGCAGCTACTGCAT
TGGGAGGGAAATCACTTATCGATCCCAAACAGGTATATGCTGGAACGGTCAAGGAACTGGAT
GAACTTCCCTTTAATATCCCGGCAGACTACAAACCGTTTACCCATCAAAGGAATATATGGGG
CCAGGCTTTATTGGGAGTACCCGAACCTCTAGCACTCAGAGAGCGTTTTGCTGAAGTAAGAT
GGAATGGTTGGCAGACAGATGGTTCGCCCGGTCTTACTGTACTTGATGGTGCGGCTGCTCAT
GCAAGCTGGGCCGTTGATTATTATCTTAACGGGGAAAATAGCGCCTGCAGGGCCAATAAAGG
TTTTTTTGAATGGCATCCCAAAGTGCCCGAGCTGAACTTTAGGTGGGGCGATCCGGAGAGAA
ATATTCATTCCCCCGGTGTAAAAAGTGCCGAAGAAGGAACGATGGCAGTAAAAAAAATAGC
TAGATTTTTCGGCGCTGCTAAAGCTGGGATAGCGCCTTTTGACAAACGTTGGGTTTTTACTGA
AACGGCTGCCTTTGTTAAAACGCCTGAGGGTGAAAGTCTGAAATTTATCCCTCCGGATTTTGG
GTTTGAGCCCAAGCATGTAATCTCGATGATTATCCCACAGTCGCTAGAAGGAATAAAGTGTG
CCCCGTCCTTTTTAGGATCAGCTGAATATGGATTAAGTTATACCCAGATTGGATATGCTGCAT
TCGGTTTATCCATGTTTATTAAAGATCTGGGATATCATGCGGTTCCAATCGGAGCTGACAGTG
CATTAGCTATGCCTATAGCTATTCAGGCGGGTCTGGGGGAATACAGCAGGTCGGGGCTAATG
ATTACGCCTGAATTTGGTCCAAATGTTAGACTCTGTGAAGTATTTACTGACATGCCTTTAAAT
CATGATAAACCTATTTCATTCGGAGTAACTGAATTTTGCAAAACCTGCAAAAAATGCGCTGA
AGCATGCGCCCCTCAAGCTATTAGCTATGAAGATCCTACCATTGATGGACCTCGTGGGCAAA
TGCAAAATTCGGGAATAAAGAGATGGTATGTTGACCCGGTGAAGTGCTTTGAATTCTGGTCG
CGTGATAACGTCAGAAACTGCTGCGGAGCTTGTATAGCTGCTTGCCCATTTACTAAG

Translation of the coding region of Clone3:
MDKEKSNNDKPATKINRRQFLKFGAGASSGIAIATAATALGGKSLIDPKQVYAGTVKELDELPFN
IPADYKPFTHQRNIWGQALLGVPEPLALRERFAEVRWNGWQTDGSPGLTVLDGAAAHASWAVD
YYLNGENSACRANKGFFEWHPKVPELNFRWGDPERNIHSPGVKSAEEGTMAVKKIARFFGAAK
AGIAPFDKRWVFTETAAFVKTPEGESLKFIPPDFGFEPKHVISMIIPQSLEGIKCAPSFLGSAEYGLS
YTQIGYAAFGLSMFIKDLGYHAVPIGADSALAMPIAIQAGLGEYSRSGLMITPEFGPNVRLCEVFT
DMPLNHDKPISFGVTEFCKTCKKCAEACAPQAISYEDPTIDGPRGQMQNSGIKRWYVDPVKCFEF
WSRDNVRNCCGACIAACPFTK

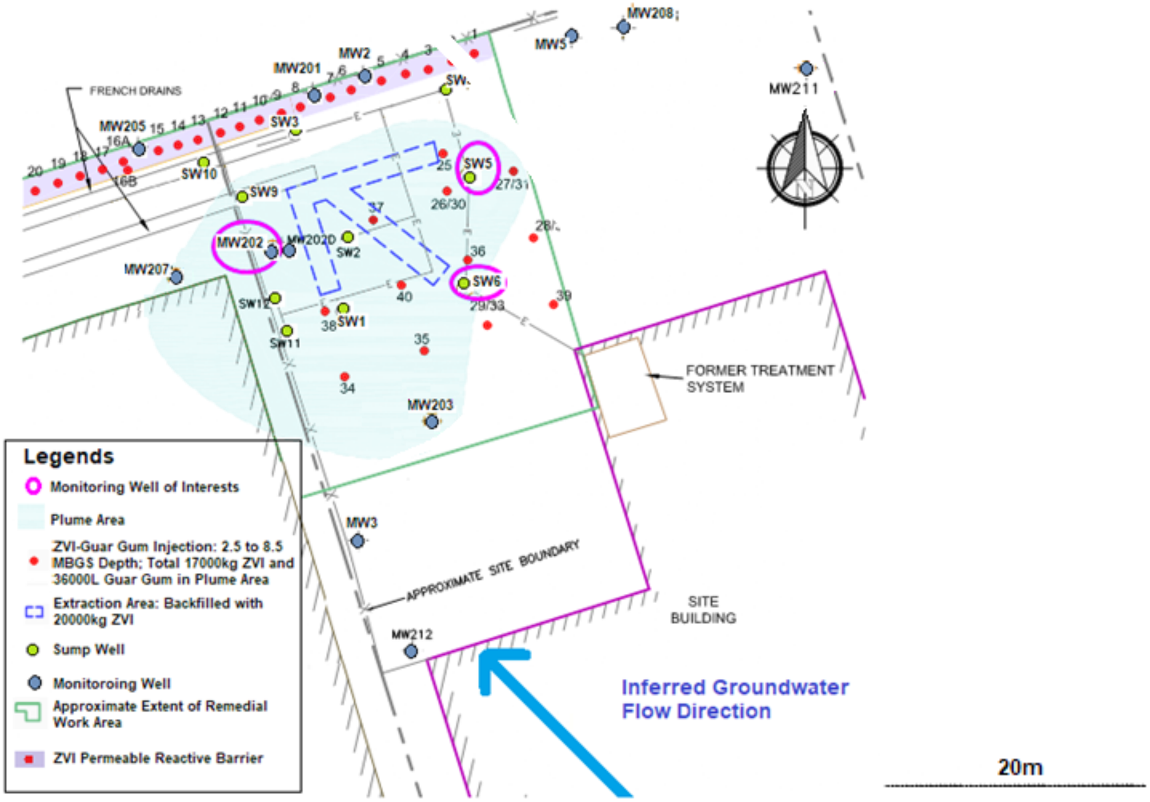

**Fig. S1. Map of the remediation site.**
Groundwater samples were collected for chemical concentration measurements and microbial analysis from the monitoring wells circled in pink (i.e., MW202, SW5 and SW6*).

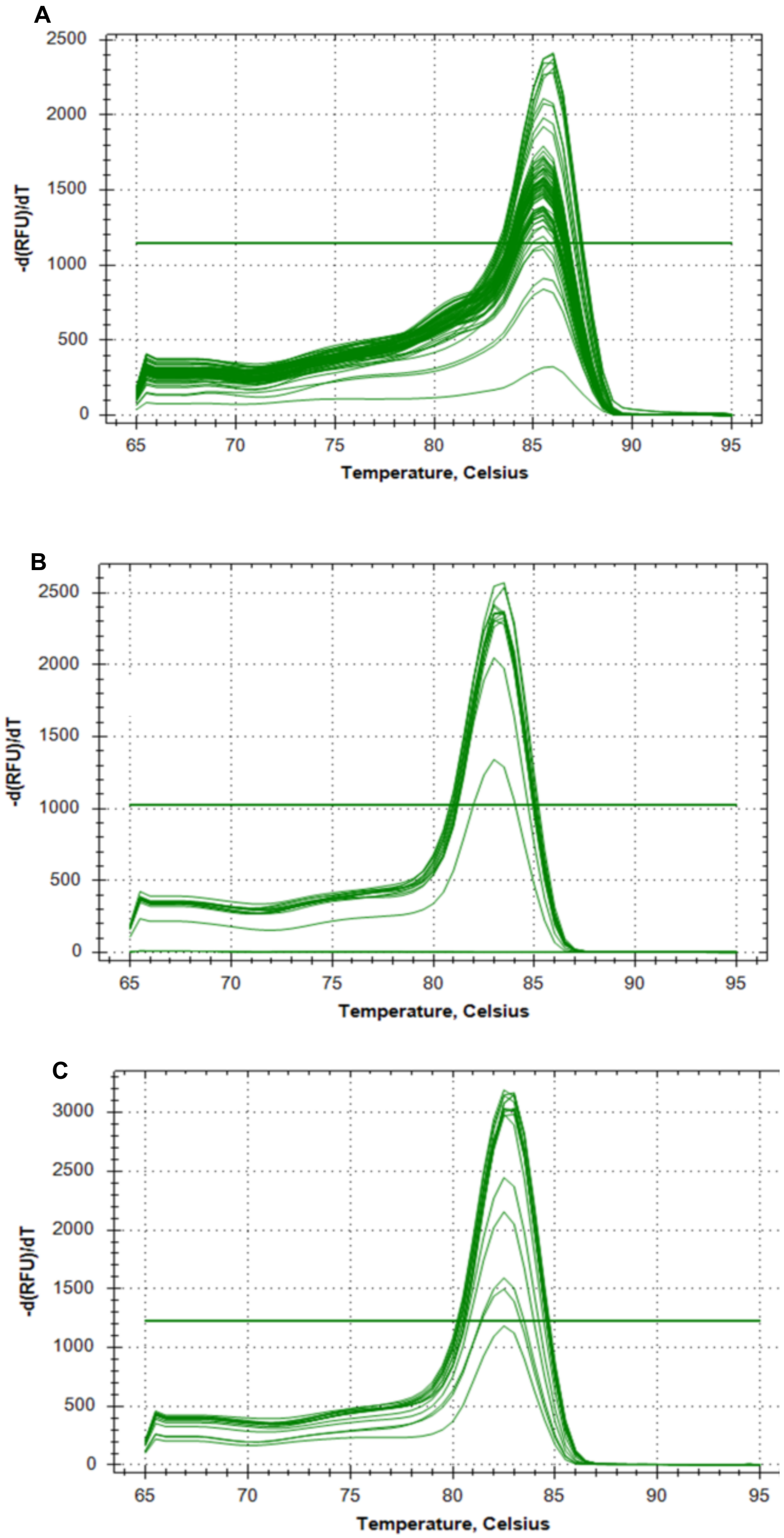

**Fig. S2. Melt curves of qPCR**

Panel A: qPCR with *Dhb* 16S rRNA primer set; Panel B: qPCR with *cfrA* primer set; Panel C: qPCR with *dcrA* primer set. Note that the products with nonspecific bindings (i.e, flagged with '*' in qPCR result figures) were excluded in these melt curves.

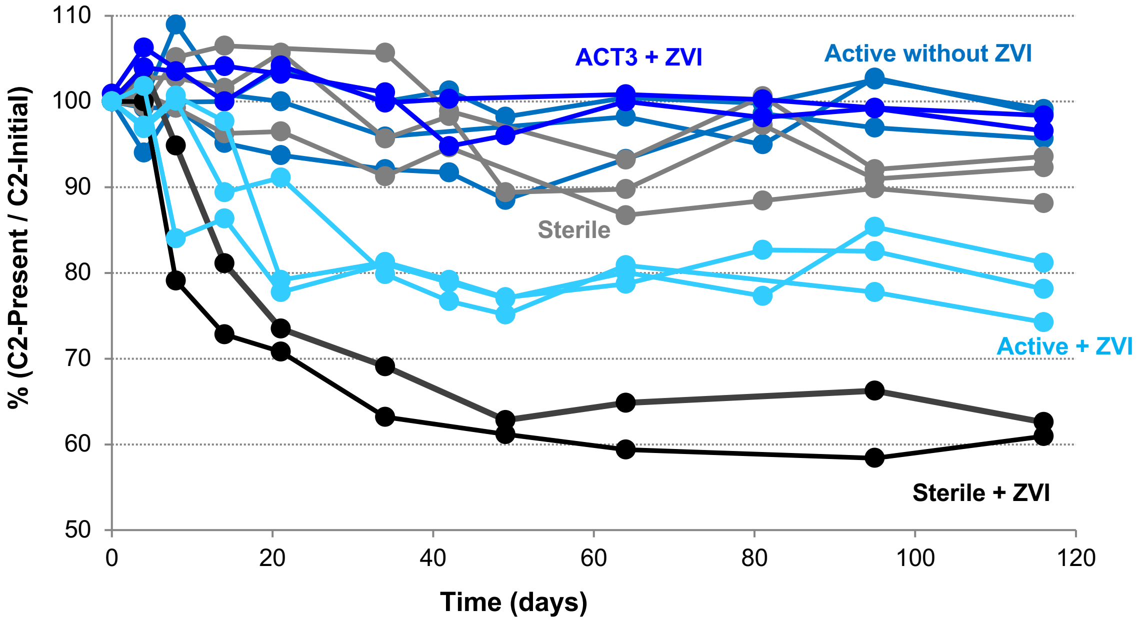

**Fig. S3. Mole balance of C2 compounds**

Data are presented as percentage of the initial concentration over the first 120 days.
Active without ZVI (teal blue), Sterile (grey) and ACT3 + ZVI (dark blue) exhibited little mole loss; Sterile + ZVI (black) and Active + ZVI (light blue) had 20-30% mole loss prior to day 30.

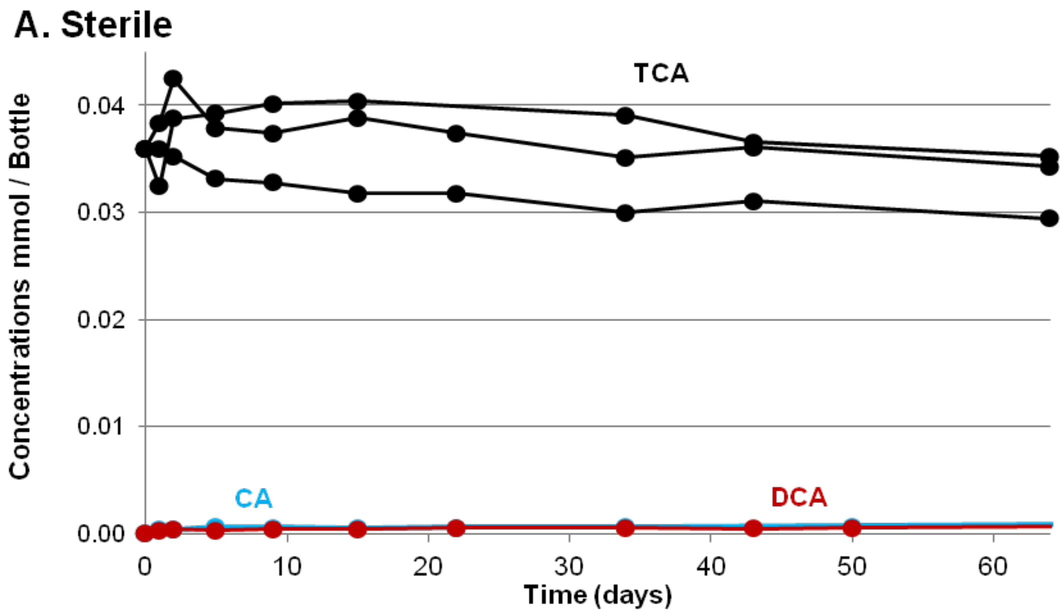

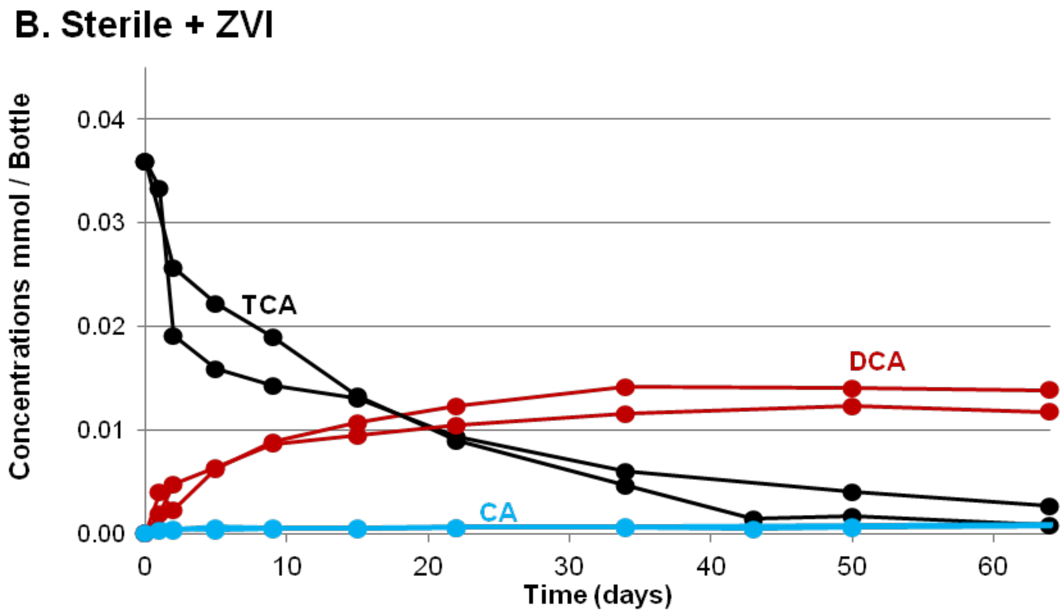

**Fig. S4. Chlorinated ethane concentration profiles in the Sterile set and in the Sterile + ZVI set**

Panel A: Sterile (negative control); Panel B: Sterile + ZVI; 1,1,1-TCA: black; 1,1-DCA: red; CA: blue.

No degradation was shown in the Sterile bottles, while 1,1,1-TCA was transformed into 1,1-DCA in the Sterile + ZVI bottles (B). No CA was generated in either case.

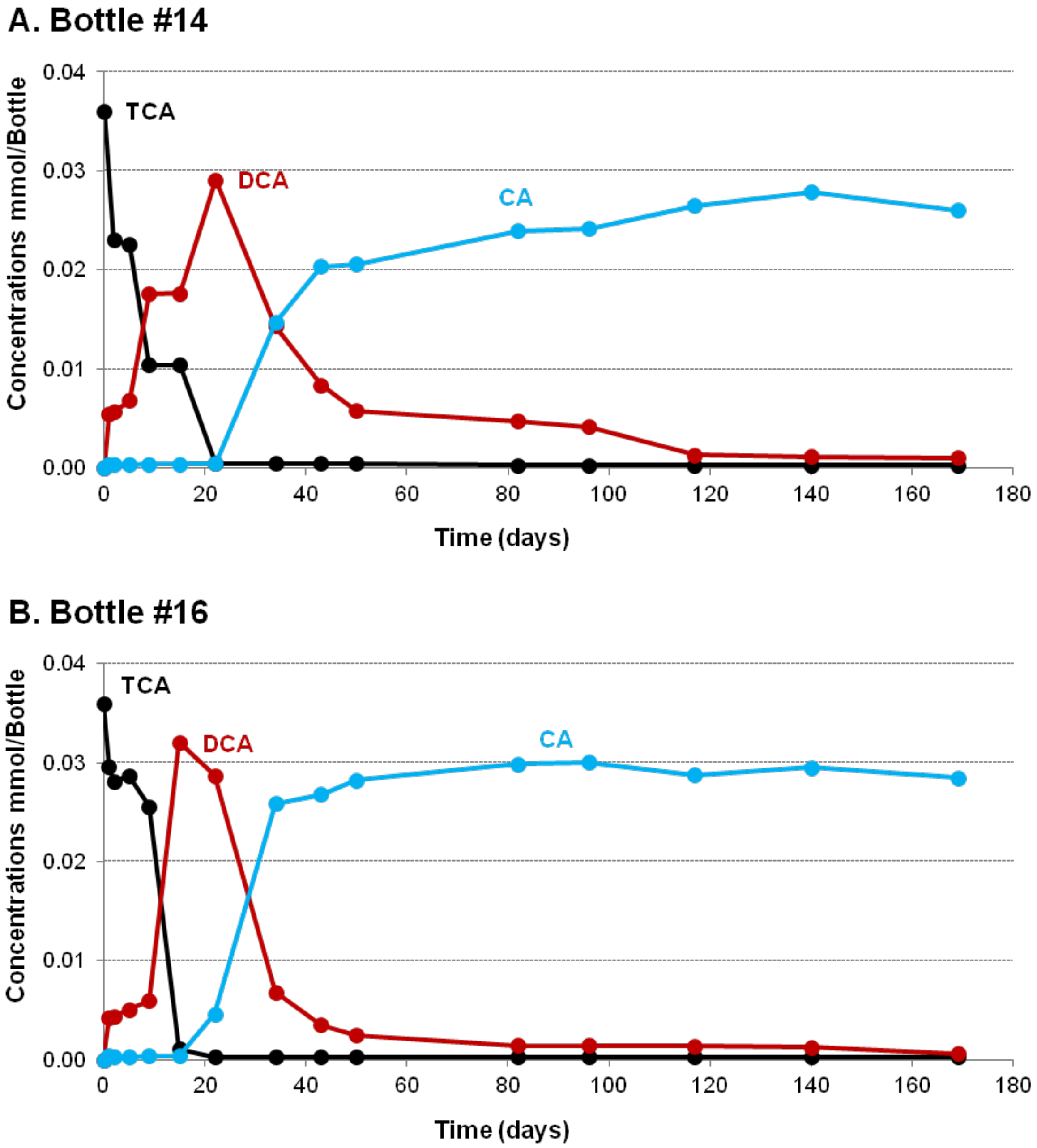

**Fig. S5. Chlorinated ethane concentration profiles in the ACT3 + ZVI bottles.**
In both bottles, 1,1,1-TCA (black) was quickly converted to 1,1-DCA (red) then to CA (blue).

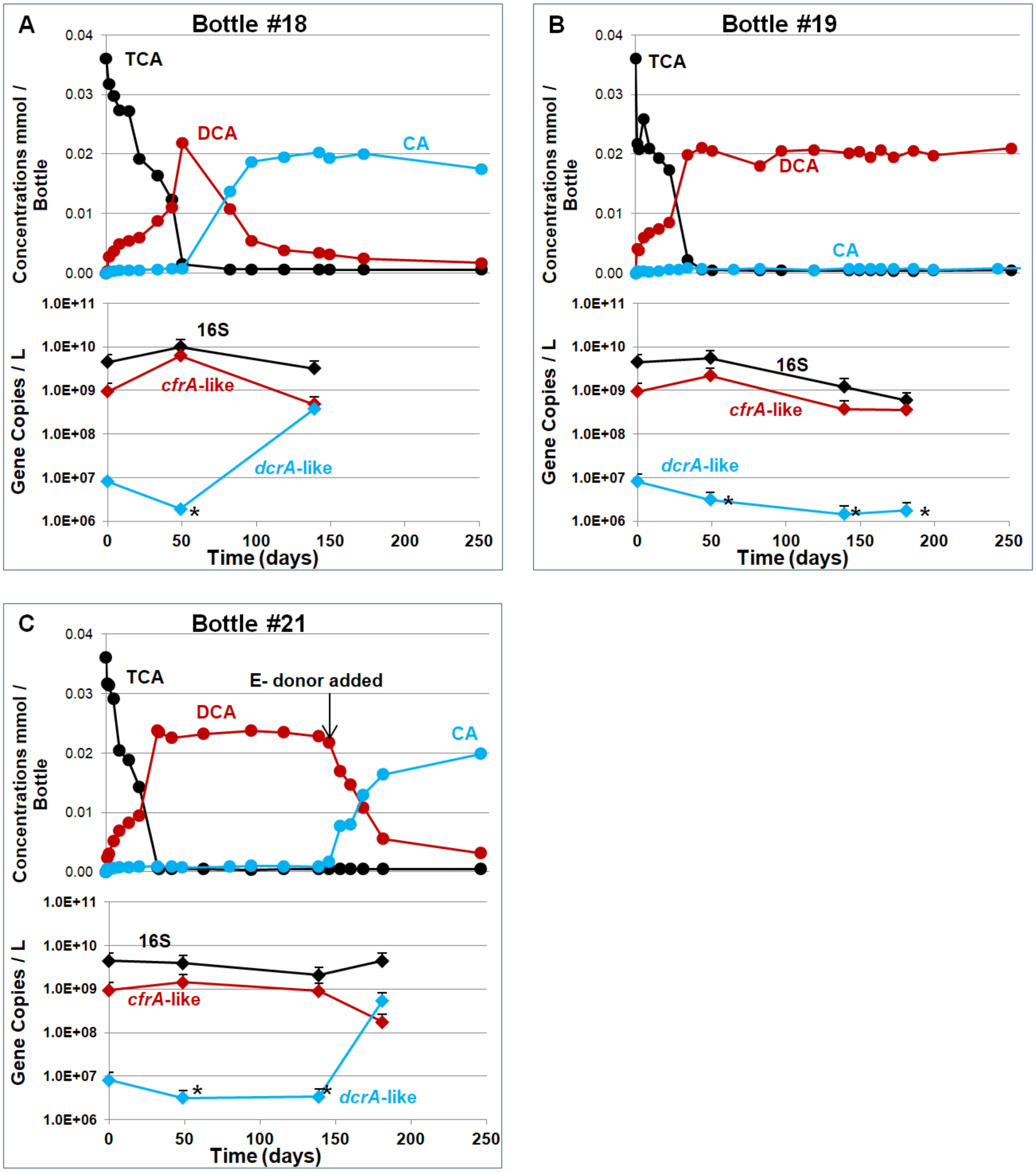

**Fig. S6. Chlorinated ethane concentrations and gene copy numbers in the groundwater microcosms with ZVI addition (Active + ZVI set) over 250 days.**

Data from bottles #18, #19 and #21 are presented in Panels A, B and C, respectively. In each panel, the top figure shows chlorinated ethane concentrations (1,1,1-TCA in black circles, 1,1-DCA in red circles, and CA in blue circles) and the bottom figure displays the qPCR results in copies per L (black diamonds for the *Dhb* 16S rRNA gene, red diamonds for the *cfrA*-like gene, and blue diamonds for the *dcrA*-like gene). Gene copies are the average of three technical replicates, and qPCR data points marked with * are below the lowest detection limit or showed nonspecific amplification.

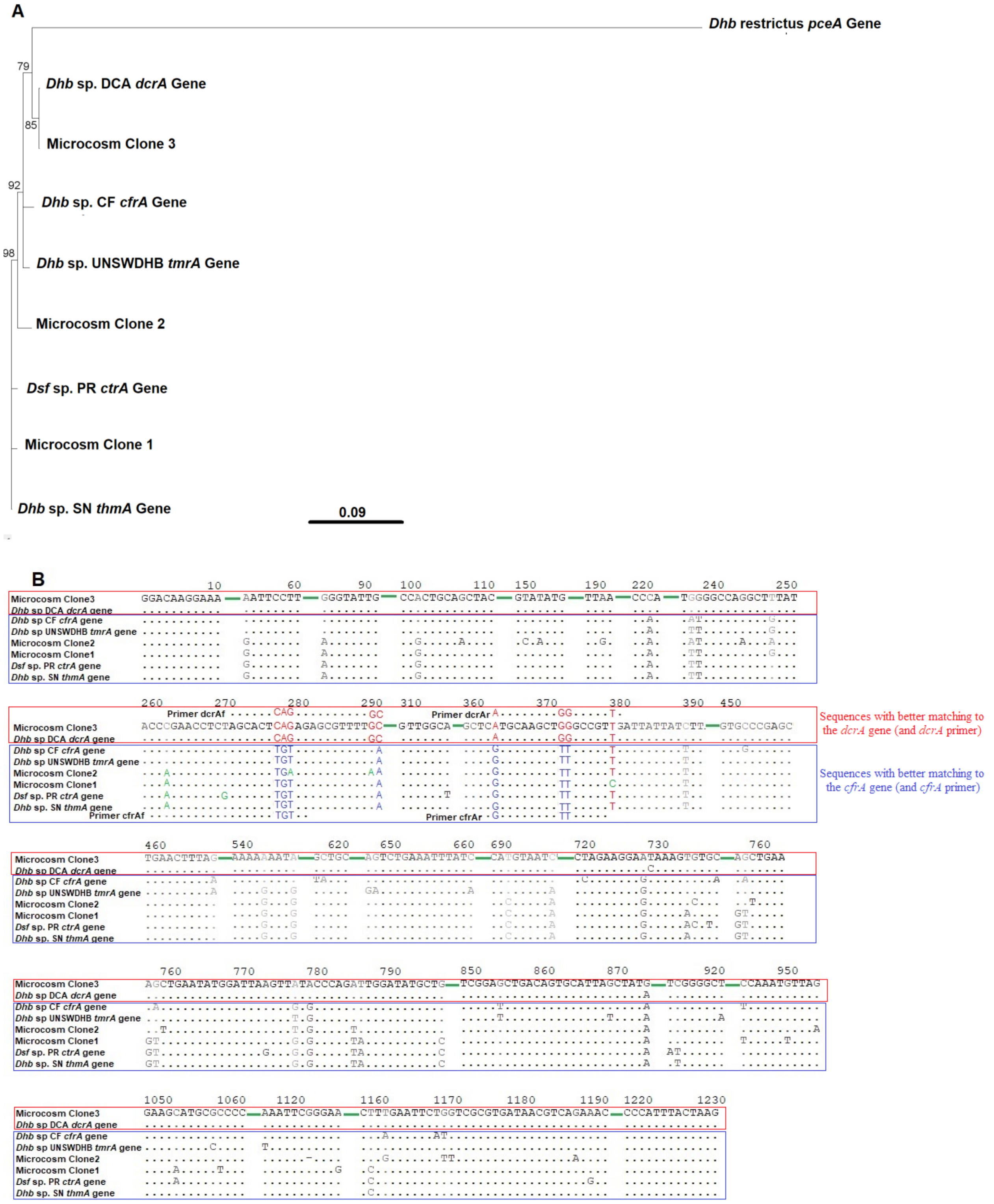

**Fig. S7. Alignment of cloned sequences to reference *rdhA* genes** **(continued next page)**

Alignment also shows location of *cfrA* and *dcrA* qPCR primers.

Panel A: tree view, with *pceA* gene encoding the tetrachloroethene reductive dehalogenase in *Dehalobacter restrictus* DSM9455 (AJ439607.2) as an outgroup to demonstrate the high similarity of the *cfrA*- and *dcrA*-like genes listed in this paper.

Panel B: alignment view indicating that the sequences fall into two clay (red box for sequences with better matching to the *dcrA* gene, blue box for sequences with better matching to the *cfrA* gene)—only regions with mismatches are shown here; the green solid lines between sections are the conserved regions; dots indicate nucleotides identical to the ones in Microcosm Clone3 sequence at the same position; the mismatches within the region targeted by the primer sets are colour coded as followed: red font shows matching to the *dcrA* primer but different than the *cfrA* primer, blue shows matching to the *cfrA* primer but different than the *dcrA* primer, and green indicates mismatch to both primer sets.

Note: the sequences of all genes and clones were trimmed down to 1236 nucleotides after sequence alignment for comparison and tree construction.

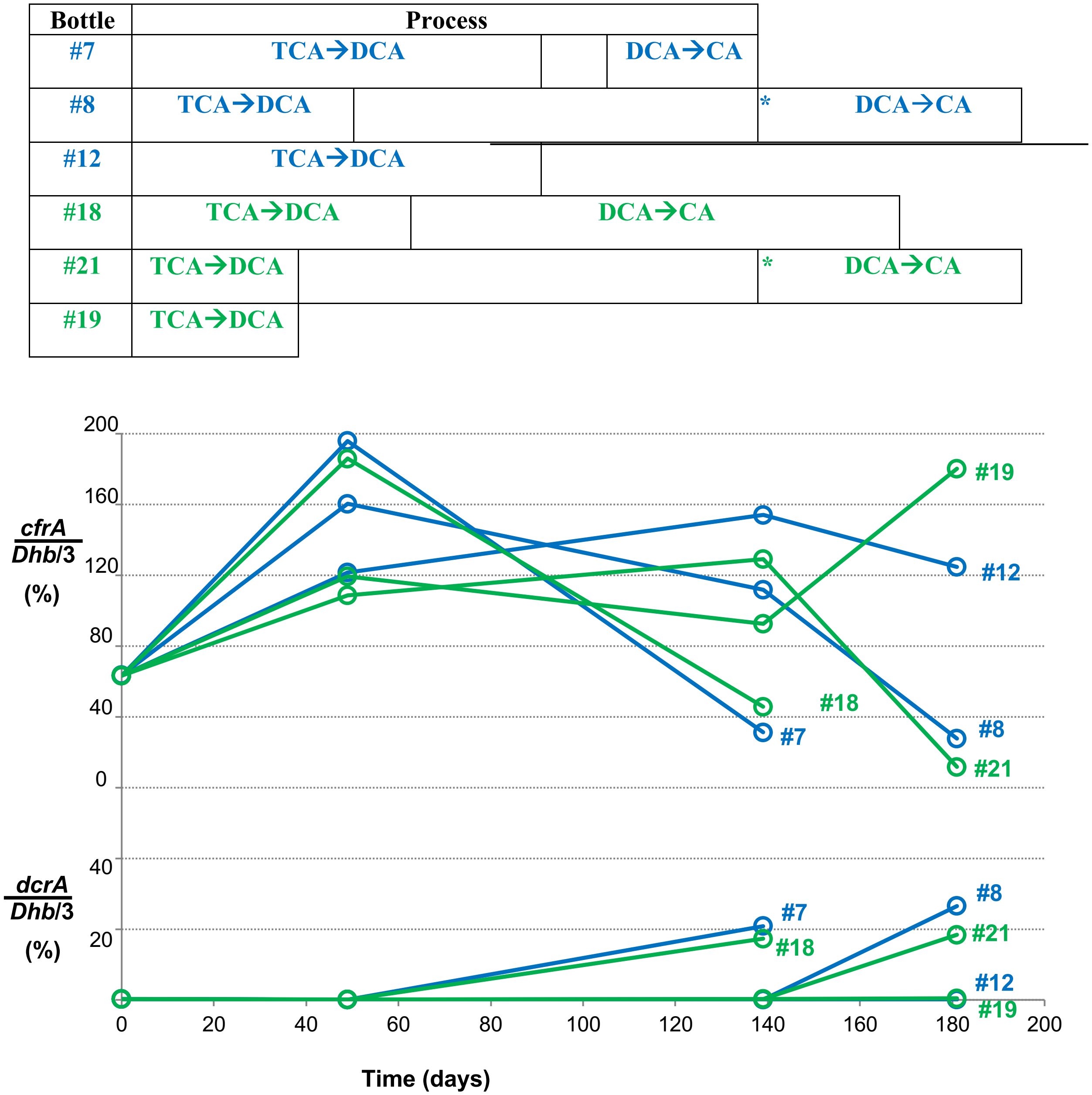

**Fig. S8. Proportion of the *Dehalobacter* population carrying the *cfrA*- or *dcrA*-like genes**

Data are from microcosms at different stages of 1,1,1-TCA dechlorination.
Active without ZVI: blue; Active + ZVI: green; Top section: illustration of degradation process; Middle section: percentages of *Dehalobacter* carrying the *cfrA*-like gene; Bottom section: *Dehalobacter* carrying the *dcrA*-like gene.

Electron donors were added to bottles #8 and #21 at the time indicated by * in the process diagram.

Note that according to the closed genomes of *Dehalobacter* sp. strain DCA (NC_018866.1) and *Dehalobacter* sp. strain CF (NC_018867.1), *Dehalobacter* usually contain three copies of the 16S rRNA genes in their genome, and only one copy of the *cfrA* or *dcrA* gene in each strain (Tang and Edwards, 2013B). Therefore, we included a factor of three when we calculated $\%_{cfrA\text{-}Dhb}$ and $\%_{dcrA\text{-}Dhb}$. As shown in the figure, the sum of these two percentages in the samples mostly fell between 75% and 150%, which is reasonable considering the errors of qPCR and that not all *Dehalobacter* contain three copies of the 16S rRNA gene (e.g., five copies in *Dehalobacter restrictus* strain PER-K23, NZ-CP007033 (Kruse et al., 2013)).

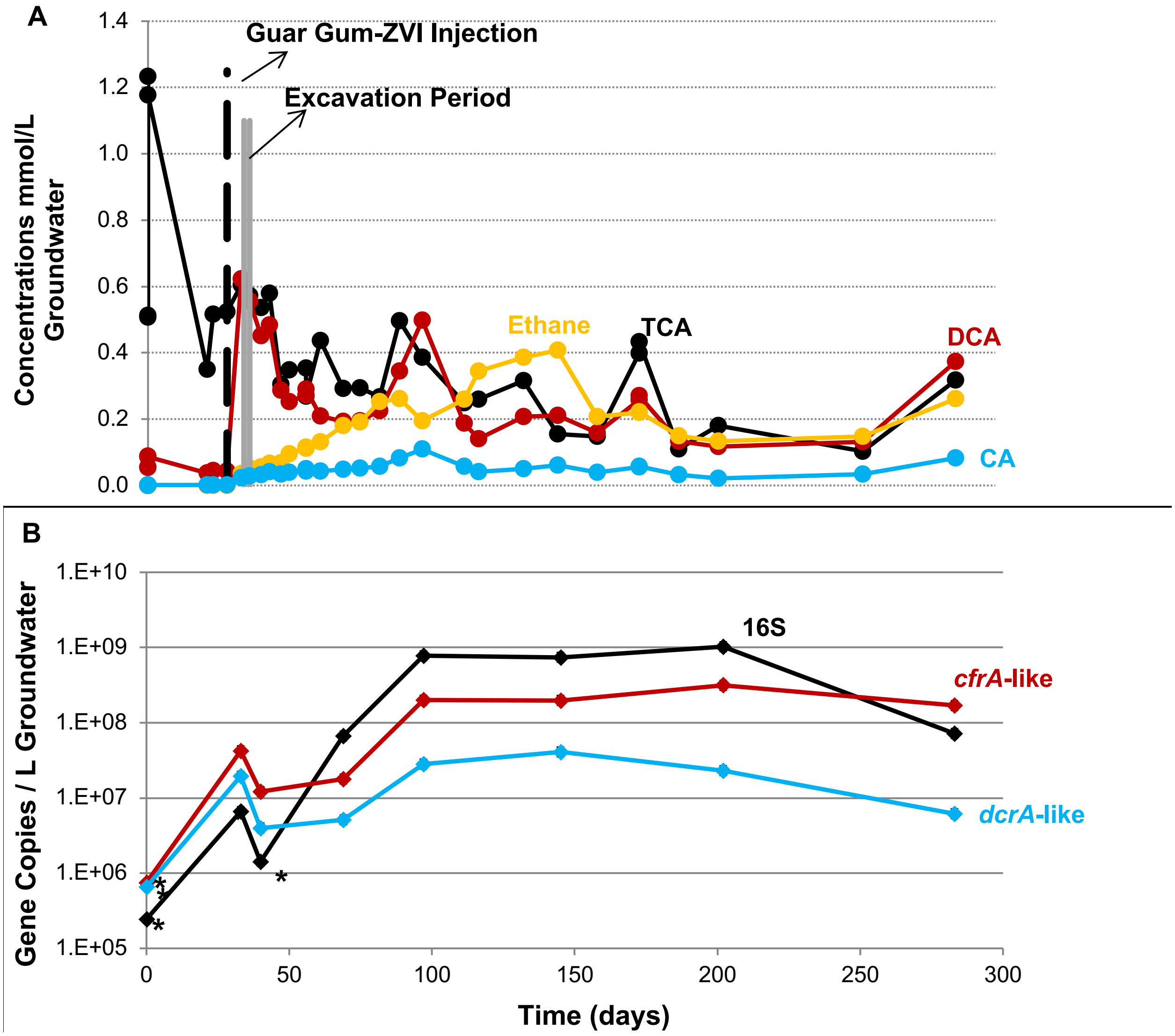

**Fig. S9. Groundwater monitoring results for MW202.**

Panel A: concentrations of 1,1,1-TCA (blank circles) and 1,1-DCA (red circles) slightly decreased after excavation followed by fluctuating chlorinated ethane concentrations. Panel B: qPCR data revealed increases in the *Dhb* 16S rRNA gene (black diamonds) and the *cfrA*-like gene (blue diamonds) after excavation upon day 96. Data points marked with'*' were below the lowest detection limit or showed nonspecific amplification products. Error bars on qPCR data reflect typical error of about a half an order of magnitude from DNA extraction and qPCR

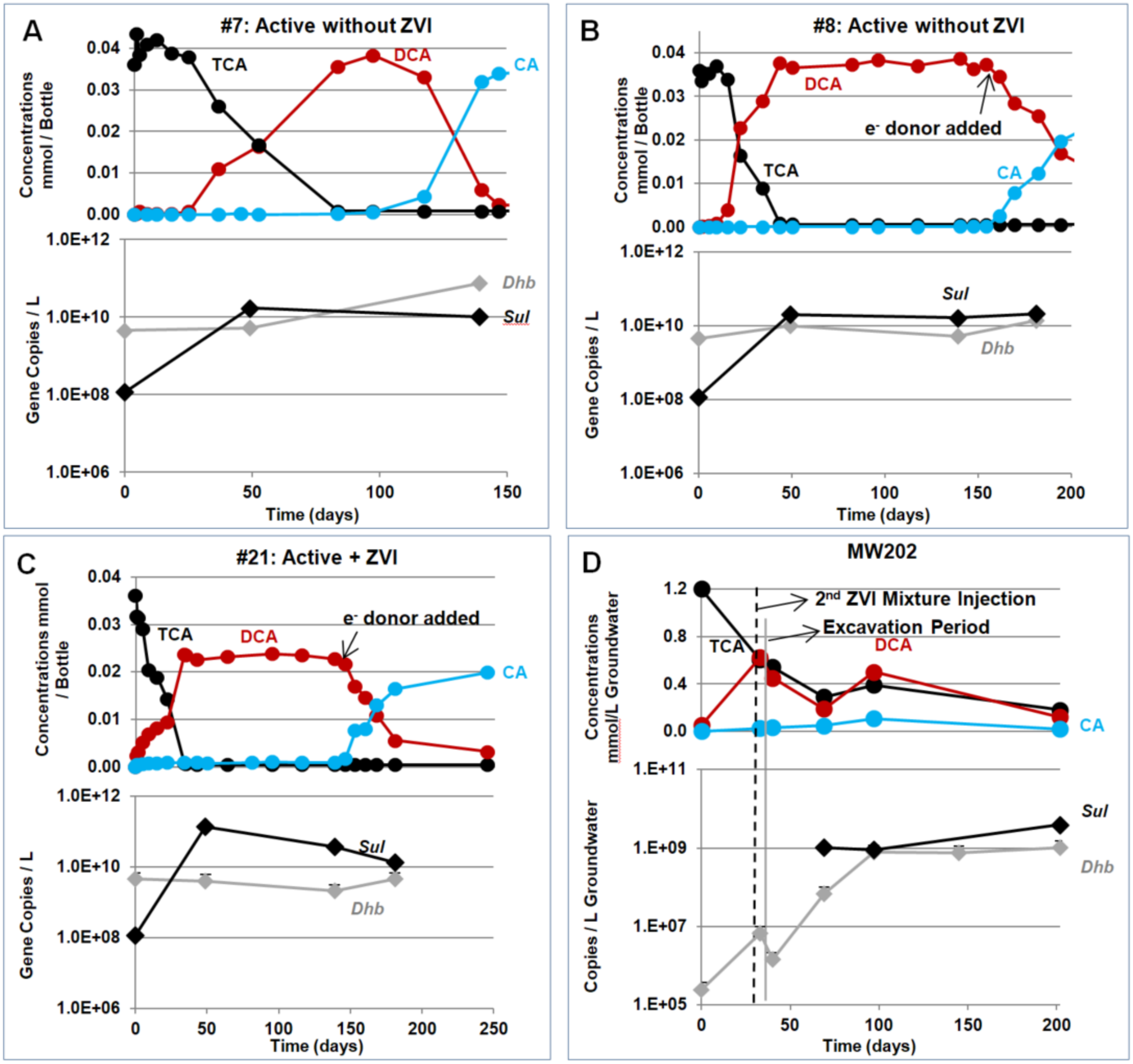

**Fig. S10. Profiles of *Sulfurospirillum* during 1,1,1-TCA and 1,1-DCA dechlorination**

Panels A to C: microcosm samples; Panels D: site samples (MW202). In each panel, the top figure shows the concentrations of chlorinated ethanes (in circles, black for 1,1,1-TCA, red for 1,1-DCA and blue for CA), and the bottom figure shows the copy numbers of the 16S rRNA gene of *Sulfurospirillum* (*Sul*, black diamonds) in comparison to the data for the 16S rRNA gene of *Dehalobacter* (*Dhb*, grey diamonds) as references. The concentrations of chlorinated ethanes and the16S *Dhb* are the same data as presented previously in Figures 1, S6 and S9. The copy number of the 16S rRNA gene of *Sulfurospirillum* (*Sul*) was estimated using the formula below:

$$Sul \text{ abundance} = Dhb \text{ abundance} \times \frac{\text{Total } \%Sul}{\text{Total } \%Dhb}$$

where *Dhb* abundance was measured using qPCR, and Total %*Dhb* and Total %*Sul* were obtained from Illumina amplicon sequencing by summing the percentages of all features belonging to *Dhb* or *Sul* respectively

Note that the field samples prior to day 69 contained 0% *Dhb*, so *Sul* abundance could not be calculated using the formula above, and thus were left blank in Fig S10.

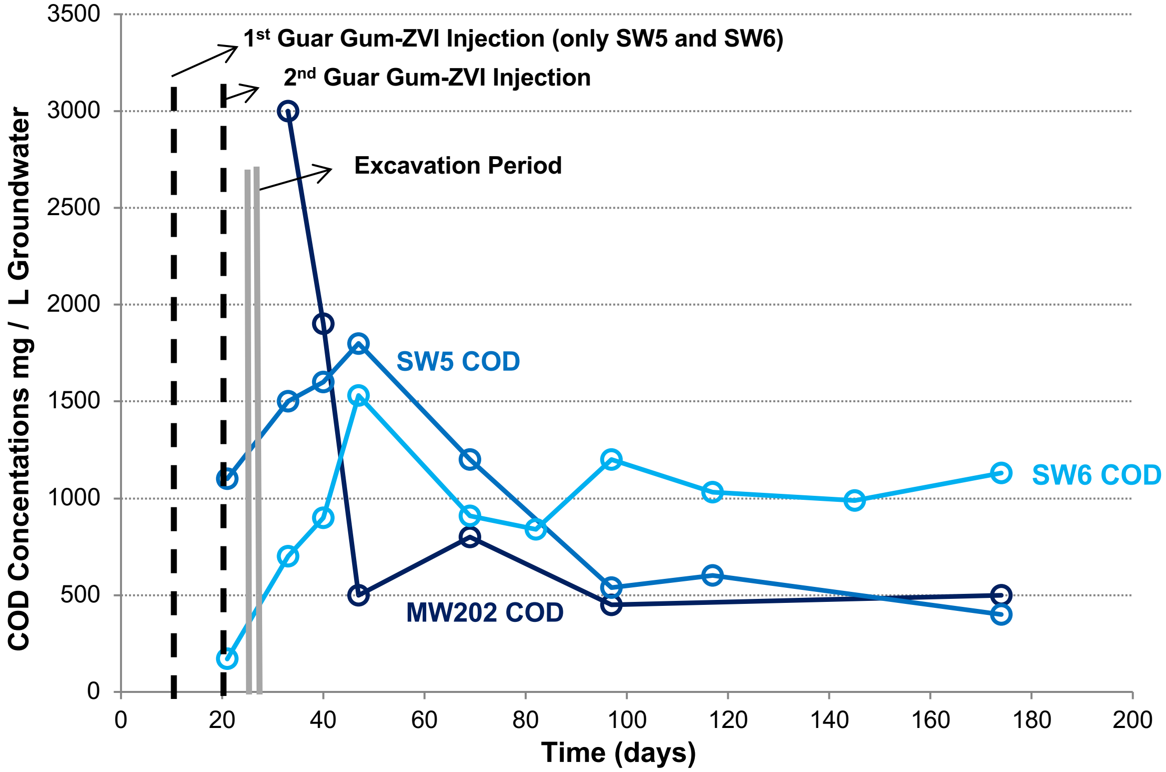

**Fig. S11. COD concentrations in groundwater samples**

SW6: light blue; SW5: blue; and MW202: dark blue
